# Huntingtin and Celsr3 Orchestrate Spontaneous Long-Distance Spinal Cord Regeneration

**DOI:** 10.64898/2026.10.01.755968

**Authors:** Jaffna Mathiaparanam, Maria F. Navarro, Jessica C. Nelson, Michael Granato

## Abstract

Spontaneous spinal cord regeneration is an endogenous process operational in diverse vertebrate and select mammalian species. Yet despite resulting in functional regeneration, axonal regrowth dynamics as well as underlying molecular programs are largely unknown. Live-cell imaging following laser axotomy of individual zebrafish spinal cord M-cell axons reveals a time-sensitive multistep process. From a genetic screen we uncover previously unrecognized roles for two evolutionary conserved genes, *celsr3* and *huntingtin* (*htt*) in spontaneous spinal cord regeneration. Specifically, we find that *htt* is critical to initiate a transcriptional injury response in regenerating M-cells, enabling their axon to traverse the injury site. Once axons cross the injury site *celsr3* functions to increase growth rates, while both *celsr3* and *htt* function to sustain long-range axonal growth. Moreover, we show that *htt* is dispensable for optic nerve and peripheral motor neuron regeneration, and that reducing soma-to-injury site distance in *htt* mutant M-cell axons partially restores regeneration, demonstrating a selective role in long-range CNS axon regeneration. Consistent with a transport-based mechanism, genetic knockdown of *htt*-associated axonal transport proteins recapitulates the *htt* mutant regeneration phenotype. Combined, our results provide compelling evidence for a time and distance-sensitive multistep molecular framework in which *celsr3* dependent growth rate modulation and *htt* dependent axonal transport promote spontaneous, long-range spinal cord regeneration.

**Significance Statement:** Unlike humans, several vertebrates — including zebrafish — as well as select mammals have retained an endogenous capacity for adult spontaneous spinal cord regeneration. First reported in rabbits by Murray et al. in 1965 [1], and more recently documented in naked mole rats and the spiny mouse (*Acomys*) [2, 3], this provides compelling evidence for evolutionarily conserved molecular programs promoting regrowth of injured spinal cord axons towards their original targets. Yet despite decades since its discovery, the identity of these molecular programs including how individual genes within programs act and interact with each other to initiate injured axons to regrowth, direct them across the injury site and sustain long-distance regrowth— has remained largely unknown.

Our study addresses this long-standing gap directly. Using live-cell imaging of laser-axotomized zebrafish spinal cord axons, we deliver the first minute-by-minute timeline of spontaneous spinal cord regeneration in a live intact vertebrate and at single-axon resolution, revealing a time- and distance-sensitive multistep process. We identify two evolutionarily conserved and previously unrecognized molecular drivers of this process and define a molecular framework in which *celsr3*-dependent growth rate modulation and *htt*-dependent axonal transport act as distinct, stage-specific regulators of spontaneous long-distance spinal cord regeneration.

## Introduction

Functional axon regeneration is a complex, multistep process. Regeneration success requires injured neurons to re-form growth cones at the proximal axon stump, initiate axon growth and navigate toward to their original targets to reestablish functional synapses. Axons of the mammalian Central Nervous System (CNS), including humans, display very limited regenerative capacity following injury, partly due to inhibitory mechanisms and non-permissive environmental substrates [1–6]. Furthermore, robust regeneration requires injured neurons to regulate intrinsic growth-promoting pathways [7–10]. For example, in adult mice, forced induction of mTOR- and Stat3-dependent pathways after injury results in increased, yet often ectopic axonal regrowth and synaptic connections, underscoring the importance to tightly control activation of neuron-intrinsic pathways [11–13]. In mammals, unaided or spontaneous CNS axon regeneration has been observed in mice, rats, and opossum at neonatal stages [14–16], as well as in select adult mammals including rabbits [17], and more recently in the spiny mouse (*Acomys*) and the naked mole rat leading to functional regeneration [18, 19]. Combined, this strongly argues for the existence of evolutionarily conserved genetic programs that have the capacity to promote robust, functional CNS regeneration. Yet despite its well documented existence, the complement of genetic factors promoting spontaneous CNS axon regeneration as well as which step within the multistep regeneration process is controlled by which factor, or combination of factors, is extremely limited.

Similarly, dynamics of CNS axon regrowth after injury has been studied extensively using cultured neurons and more recently with three-dimensional neural tissue models [20, 21]. While these models enable precise studies at near-single-axon resolution, quantifying the dynamic behaviors of regenerating vertebrate CNS axons in their natural environment early during the regeneration process still poses a challenge. For example, whether axonal regrowth rates are fixed or tunable, e.g. whether axons navigating the injury site grow with the same growth rates and dynamics as they do when approaching their original synaptic targets is currently unclear. We previously established the larval zebrafish Mauthner cell neurons (M-cell) as a vertebrate model to visualize, quantify and genetically dissect spinal cord regeneration *in vivo* [22] . The zebrafish M-cells are a pair of giant bilateral reticulospinal hindbrain neurons, and essential components of the circuitry mediating the acoustic startle response [23–25]. Each M-cell projects a single, large diameter and myelinated axon that extends from the hindbrain to the distal tip of the spinal cord, representing the longest and largest diameter zebrafish CNS axon. Following laser axotomy, M-cell axons robustly regenerate to pre-injury length within 96 hours of injury, with functional circuit recovery evident 8-10 days after injury [22, 26]. As part of an ongoing genetic screen for genes that control eary steps of long-range M-cell axon regeneration, we identified the *Cadherin EGF LAG seven-pass G-type receptor 3* (*celsr3*) gene, best known for its role in planar cell polarity [27, 28], and the *huntingtin* (*htt*) gene to regulate long-range M-cell spinal cord regeneration. Live cell imaging reveals that in wild-type animals, regenerating M-cells axons, after crossing the injury site abruptly increase growth rates by almost 3-fold. We find that *celsr3* is required selectively for this switch in extension rate, as mutants fail to increase axonal growth rates, resulting in stunted axon regrowth. In contrast to *celsr3* predominantly regulating a single step, we find that *htt* operates at multiple steps during the regeneration process, first in initiating growth cone morphology and motility to rapidly traverse the injury site, and later to sustain long-range axonal growth. Moreover, we find that *htt* is critical for the timely onset of a transcriptional injury response in regenerating M-cell axons, consistent with the existence of critical time windows that link transcriptional activation in injured CNS axons to their regenerative capacity. Finally, we show that the endogenous *htt* gene functions as a key regulator of distance-dependent regeneration and provide genetic evidence for *htt* dependent mechanisms of axonal transport required *in vivo* for long-range M-cell axon regeneration. Combined, our results provide compelling evidence that spontaneous regeneration of spinal cord axons is a multistep process for which we identify two evolutionarily conserved genes that individually, and in combination, control distinct steps critical for long-range axonal regrowth. More broadly our results provide a conceptual and mechanistic link between time, soma-to-injury site distance, and regenerative capacity in spinal cord regeneration.

## Results

### Spontaneous regeneration of M-cell axons is a genetically controlled multistep process

We previously established a quantifiable assay for spontaneous spinal cord regeneration focusing on myelinated Mauthner cell axons (M-cell axons) within the larval zebrafish spinal cord [22]. The regenerative potential of M-cell axons varies by injury context [26, 29], and we find that laser transection of the M-cell axons around 780 µm distal to the cell soma (Figure 1a.) results in a highly reproducible regenerative response that includes Wallerian degeneration of the now separated distal axon fragment [22]. While Wallerian degeneration of CNS axons, including its dynamism, has been studied extensively in nonmammalian and mammalian systems [30–32], the early steps and dynamic behaviors of regenerating spinal cord axons in their native environment have not been examined at single axon resolution. To quantify and classify the earliest steps of M-cell axonal regrowth emerging from the proximal axon stump following injury, we used laser axotomy and live cell imaging of GFP labeled M-cells via the *Tg(Tol-056)* transgene [33] as previously described [22]. Figure 1b is a schematic representation of morphologically distinct steps during the regeneration process from the time of axotomy until ∼ 24 hours post transection (hpt), as detailed below. We find that almost instantly following axotomy the distal and proximal tips of the severed M-cell axon fully separate and retract creating an injury gap. The distal axon fragment undergoes Wallerian degeneration between 15-48 hpt. The proximal axon stump changes its morphology within minutes of separating from the distal segment, and during the first 10 hpt, undergoes extensive remodeling, dynamically extending and retracting small transient processes into the injury gap (Figure 1b, c; Supplemental Movie 1). This transient extension and retraction step is followed by stabilization of a growth cone like tip that becomes motile, resulting in consistent forward extension of the regenerating axon into the injury gap (Fig. 1b, d; Supplemental Movie 2). By 24 hpt, regenerating M-cell axons have crossed the injury gap and frequently form multiple, caudally directed branches (Figure 1b; white arrow heads in Figure 1e). By 48 hpt, axon length has greatly increased (Figure 1f). This dramatic increase in axon length between 24 and 48 hpt correlates with a temporarily restricted, ∼ 3-fold increase in axonal growth rate (Figure 1g). After 48 hpt, growth velocities remain constant, ectopic axon branches have retracted, and by 96 hpt over 90% of regenerating M-cell axons have extended at least 70% of their pre-injury length (Figure 1h). By 120 hpt M-cell axons have extended to 100% of their pre-injury length [22] and by 8 days post injury M-cells have regained pre-injury functionality [26]. Thus, the initial steps of spontaneous regeneration of spinal cord M-cell axons occur in a stepwise fashion and are characterized by dynamic, morphological changes at the axon tip navigating the injury gap, followed by a rapid and focused acceleration in growth rates once the regenerating axon has traversed the injury gap.

**Figure 1:**
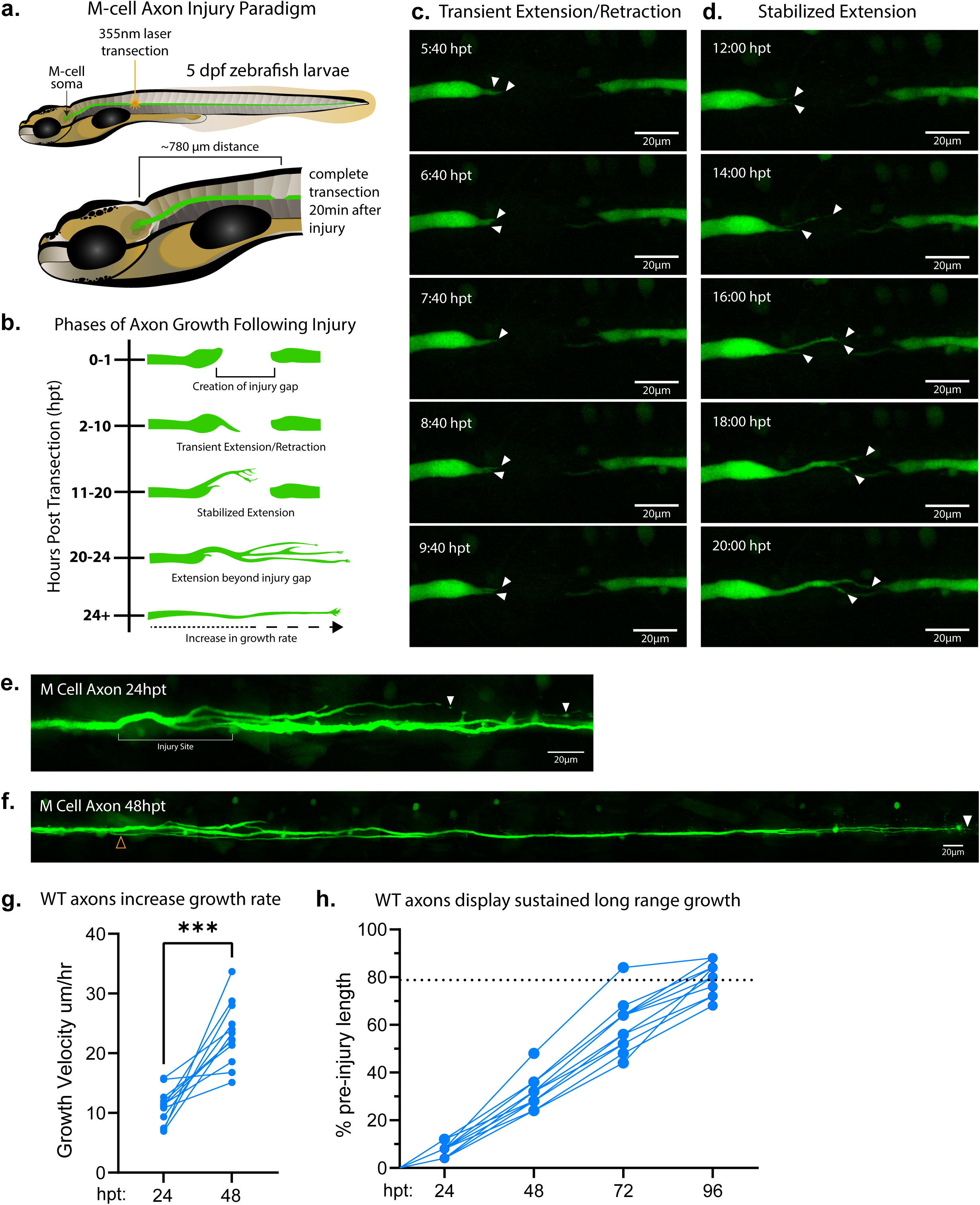
M-cell regeneration is a multistep process. (a) Schematic of laser-induced transection of M-cell axon in the zebrafish spinal cord at 5 dpf. Injury occurs approximately 780 µm from the M-cell soma. (b) Schematic of wildtype M-Cell axon growth after injury. (c) Timelapse images depicting the proximal axon undergoing a brief period of transient extension/retraction. (d) Timelapse images depicting a stabilized axon crossing the injury gap. (e) Image of a wildtype axon at 24 hpt. White arrows delineate regenerating axon tips. (f) Image of wildtype axon at 48 hpt. Yellow arrow depicts location of original axon injury, white arrow delineates regenerating axon tip. (g). Quantification of growth velocity increase in wildtype axons monitored between 0-24hpt and 24-48hpt. Wildtype axons consistently display increased growth velocity between 24-48 hpt compared to 0-24hpt (n=12). Difference in velocity measured using paired t-test (***p= 0.0001). (h) Quantification of wildtype axon regrowth at 24h intervals from 0-96hpt. Dotted line located at average percentage of pre-injury length recovered by 96 hpt (78.9 ± 6.7% pre-injury length, n=14). dpf: days post fertilization, hpt: hours post transection.

### *htt* and *celsr3* promote *in vivo* M-cell axon regeneration

In a genetic shelf screen (for details see Bremer et al 2019) we previously identified several genes required for short-distance M-cell axon regeneration by screening for axonal regrowth length at 48 hpt when wild-type M-cell axons have reached ∼50% pre-injury length. To identify genes that promote long-range M-cell regeneration, we re-screened mutants from this collection as well as additional mutant lines for axonal regeneration length at 96 hpt when wild-type M-cell axons have regrown on average to ∼79% preinjury length. From this latest screen, which include mutants of 23 candidate genes (Supplemental Table 1), we identified mutants in two genes, *celsr3* and *huntingtin* as *in vivo* regulators of long-range M-cell regeneration. *Celsr3* is a member of the adhesion GPCR gene family, encoding a G-Protein coupled receptor (GPCR) with a cadherin EGF LAG-repeat 7-transmembrane domain which functions as a co-receptor for the *wnt* receptor *frizzled* [27]. *Celsr3* is considered a core component of the Wnt/planar cell polarity (PCP) pathway which plays well-documented roles in invertebrates and vertebrate development, including in neural development [28, 34, 35]. In addition, Celsr3 has been shown to play roles in mammalian CNS synapse formation [36] and behavior [37]. *Huntingtin (htt)* encodes a 350 kDa scaffolding protein with over 300 identified binding partners. Through interactions with several transcriptional activators and repressors, *htt* has been implicated in gene regulation [38], including with the transcriptional regulator *Tumor suppressor protein 53* (*tp53*) [39, 40]. In addition, *htt* has been shown to regulate axonal trafficking through direct regulation of motor proteins [41], and proteins involved in vesicle sorting and endocytosis [42–46]. There is emerging evidence that *htt* plays a role in neural plasticity after CNS axon injury [47] [48, 49], however, *in vivo* roles of the endogenous, wild type HTT protein during the process of spinal cord regeneration as well as the *in vivo* relevant interactors through which *htt* might promote CNS axon regeneration are largely undefined. Importantly, both the *celsr3* and *htt* mutants we use in this study are caused by premature stop codons that truncate the open reading frame and likely represent null alleles (Figure 2a).

**Figure 2:**
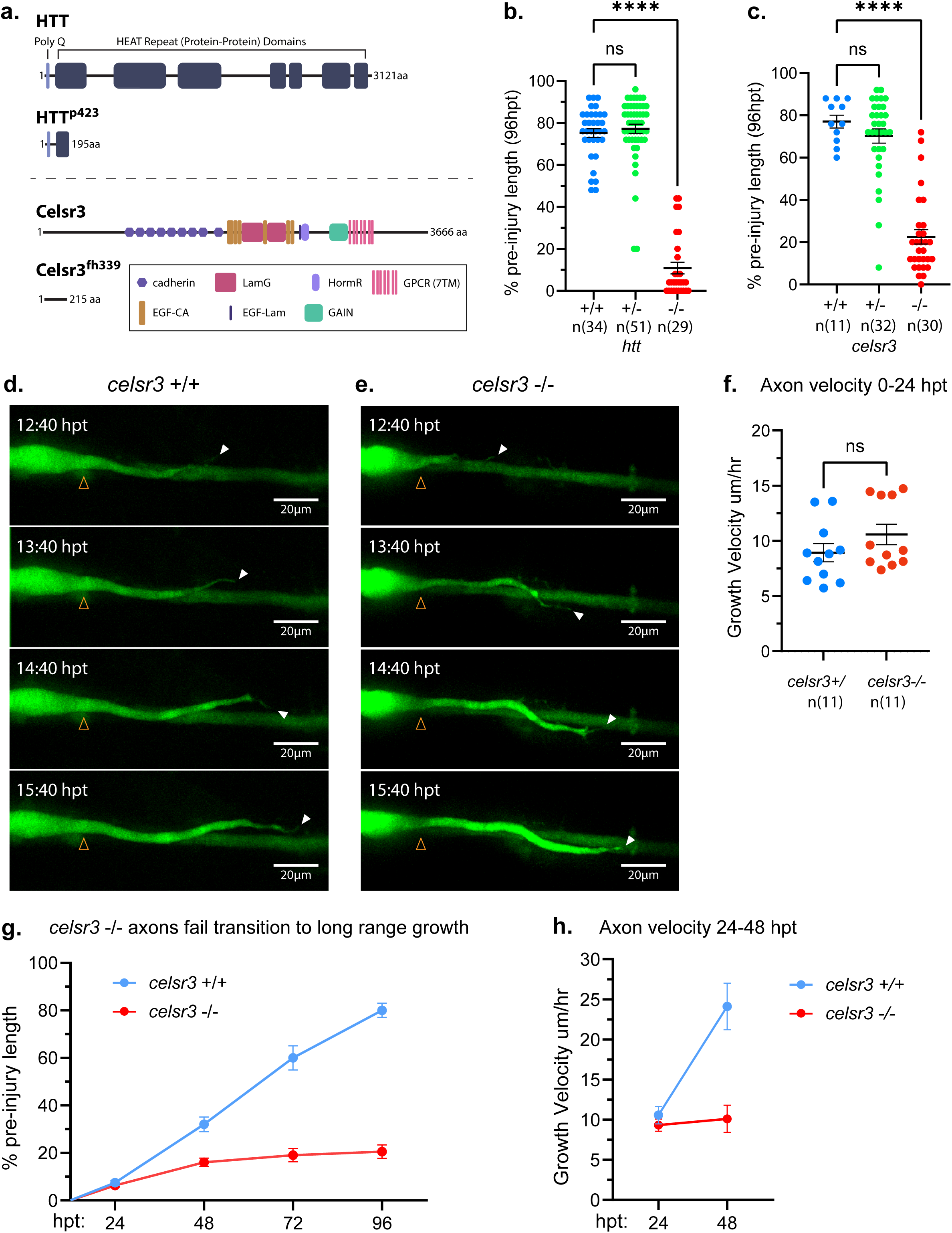
*Htt* and *celsr3* promote M-cell axon regeneration. (a) Schematic of HTT and Celsr3 with relevant protein-protein binding domains and predicted outcome of mutations. 7bp deletion in *htt* resulting in a frameshift mutation and premature stop at 197aa (allele: *htt* p423). Point mutation in *celsr3* resulting in premature stop at 215aa (allele: *celsr3* fh399). (b) Quantification of axon regeneration length at 96hpt of *htt*+/+ (75.16 ± 12.29% pre-injury length, n=34), *htt*+/- (77.16 ± 15.46% pre-injury length, n=51), and *htt*-/- (10.90 ± 14.85% pre-injury length, n=29). *Htt* mutants display significant deficits in regrowth compared to wildtype siblings (****p< 0.0001, Dunnett’s multiple comparisons test). (c) Quantification of axon regeneration length at 96hpt of *celsr3*+/+ (77.08±10.13 %pre-injury length, n=11), *celsr3*+/- (70.24±19.09 %pre-injury length, n=32), and *celsr3*-/- (55.53±18.63%pre-injury length, n=30). *Celsr3* mutants display significant deficits in regrowth compared to wildtype siblings (****p< 0.0001, Dunnett’s multiple comparisons test). (d) Timelapse images of *celsr3* +/+ sibling M-cell axon extension through injury gap. (e) Timelapse images of *celsr3* -/- mutant M-cell axon extension through injury gap. Yellow arrows depict start of regrowth; white arrows depict longest point of extension. (f) Quantification of growth cone velocity in *celsr3* +/+ (8.92 ± 2.72 µm/hr, n=11) and *celsr3* -/- (10.58 ± 3.07 µm/hr, n=11) axons imaged between 0-24 hpt. No significant difference in growth cone velocity observed during this time interval (p=0.1964, Welch’s t-test). (g) Quantification of *celsr3* +/+ and *celsr3* -/- axon regrowth at 24h intervals from 0-96hpt. (h) Relative increase in growth cone velocity between 24 and 48hpt in *celsr3* +/+, +/- and *celsr3* -/-. *Celsr3* +/+, +/- siblings have an average growth rate of 10.57 ± 2.88 µm/hr at 24 hpt that increases to 24.12 ± 7.67 µm/hr at 48 hpt (n=7). *Celsr3* -/- mutants extend at an average rate of 9.33 ± 3.04 µm/hr at 24 hpt and remain at a similar rate of 10.11 ± 6.79 µm/hr at 48 hpt (n=16). +/ :sample includes both +/+ and +/- genotypes.

To characterize the *in vivo* roles of both genes in spinal cord regeneration, we first performed static end point analyses of M-cell axon regrowth at 96 hpt. Compared to their wild-type siblings, neither *celsr3* nor *htt* heterozygotes (+/-) displayed a significant reduction in axonal regrowth length (Figure 2b, c). In contrast, homozygous mutant *celsr3* and *htt* animals display highly significant reductions in axonal regrowth length. Specifically, compared to their wild-type siblings where the majority of transected M-cell axons regrew to over 70% pre-injury length (100 out of 128 axons, pooled), in *celsr3* homozygous mutants, only 3 out of 30 transected axons regrew to 60% pre-injury length while the majority of M-cell axons (24 out of 30 axons) regrew to less than 30% of their pre-injury length (Figure 2b). Similarly, in *htt* homozygous mutant animals the majority of M-cell axons (22 out of 29 axons) regrew to less than 20% of their pre-injury length (Figure 2b). Thus, both *celsr3* and *htt* are required *in vivo* for M-cell axon long-range regeneration.

### *Celsr3* increases M-cell axonal regrowth speed once they traversed the injury gap

Our endpoint analysis revealed that the majority of *celsr3* mutant M-cell axons regrew to less than 30% of their pre-injury length (Figure 2c), suggesting that *celsr3* might regulate overall axonal regrowth rates. Alternatively, *celsr3* might promote growth selectively during a specific time period, such as during or after traversing the injury gap. To distinguish between these possibilities, we used live-cell imaging and compared growth rates of regenerating M-cell axons between wild-type and *celsr3* mutants as they initiate growth and traverse the injury gap and after they had extended beyond the injury gap. We found that M-cell axons in *celsr3* ^-/-^ animals cross the injury gap on average at a speed of 9.8 µm/hour, indistinguishable from the average we observe in *celsr3* siblings ^(celsr3+/+; celsr3+/-)^ (Figure 2d-f). In contrast, when we examined the average speed between 24 and 48 hpt, after regenerating M-cell axons had crossed the injury site, we find that unlike wild-type M-cell axons *celsr3* ^-/-^ axons fail to increase their speed to 25 µm/hour and instead extend at an injury-site crossing speed of ∼10 µm/hour (Figure 2g). Concomitantly with the failure to increase axonal growth rates after crossing, 77% (n=9) of regenerating *celsr3* ^-/-^ M-cell axons we traced between 24 and 96 failed to significantly increase in length after 48 hpt (Figure 2h). This contrasts with *celsr3* ^+/+,^ siblings in which 100% (n=7) increased regrowth length between 48-96hpt from an average of 32% pre-injury length to 70% pre-injury length. Thus, *celsr3* is dispensable for initial growth cone motility as regenerating axons navigate the injury gap, and instead is required for M-cell axons once they have successfully traversed the injury gap to almost triple their speed and engage in long-range axon regeneration.

### *Huntingtin* is required for multiple steps of M-cell axon regeneration

While the etiology of pathogenic *htt* isoforms has been studied extensively in adult and development-stage animal models [50, 51], the early embryonic lethality observed in *htt* knock out mice [52, 53] has limited the number of studies focusing on developmental and post-developmental functions of the endogenous *htt* gene. Moreover, genetic loss of function alleles in zebrafish *htt* have not been analyzed in the context of axon regeneration. We therefore first asked whether *htt* plays a role in M-cell axon development. We failed to detect anatomical or morphological deficits in *htt*^-/-^ mutants we generated (for details see Material and Methods) between 1 and 5 dpf. Analysis of M-cell development at 5 dpf revealed that *htt* ^-/-^ M-Cell axons (2216 ± 78.06 µm; n=7) are comparable in length to their wild-type siblings (2245 ± 93.44 µm; n=8; Figure 3a). Our static endpoint analyses of 96 hpt M-cell regeneration in *htt* mutants suggests the existence of multiple distinct phenotypic populations. Specifically, for 62% (n=18) of *htt ^-/-^* M-cell axons we failed to detect any significant regrowth, for 10% (n=3) we observe between 5-10% regrowth length, and 28% (n=8) of axons extended beyond 10% and up to 50% of pre-injury length (Figure 2b), suggesting that *htt* plays roles at different steps of M-cell axon regeneration.

**Figure 3:**
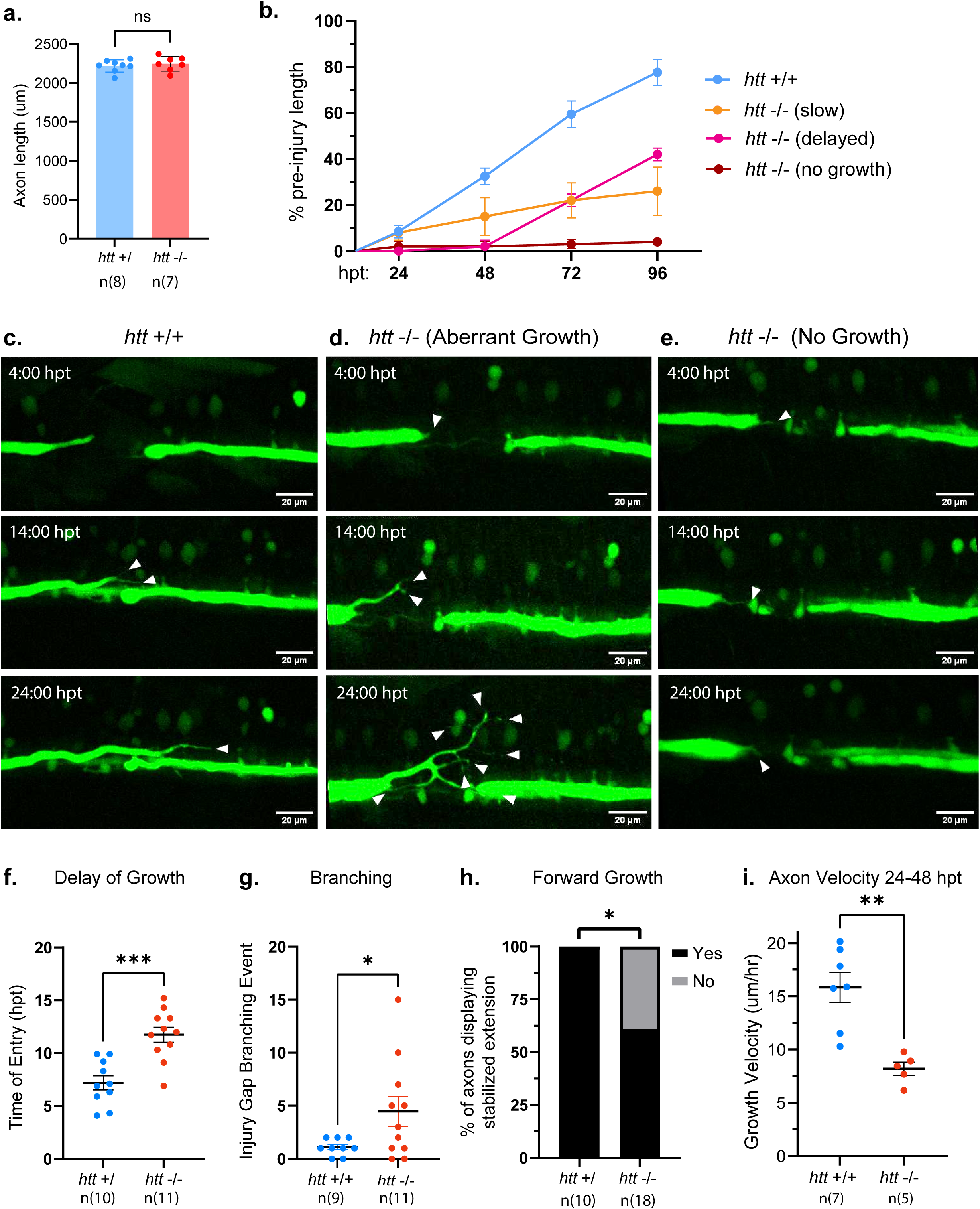
*huntingtin* is required for multiple steps in M-cell regeneration. (a) Quantification of pre-injury M-cell axon length in *htt* siblings (n=8 axons) and *htt* mutants (n=7 axons) at 5 dpf from site of transection (p=0.5326, Welch’s t-test). (b) Quantification of axon regrowth at 24h timepoints from 24-96 hpt of *htt* siblings (n=7 axons) and *htt* mutants (n=10). *Htt* mutant regrowth was defined by growth rate patterns and separated into categories of no growth (n=4/10 axons, red), slow growth (n=4/10 axons, orange), and delayed growth (n=2/10 axons, magenta). (c) Selected timelapse images of wildtype M-cell axon regrowth between 0-24 hpt. Wildtype axons display efficient growth through the injury gap with minimal branching. Axons labeled with Tol-056: GFP. (d-e) Selected timelapse images of *htt* mutant M-cell axon regrowth between 4-24 hpt. A subset of axons display increased and complex branching while navigating the injury gap. 38.8% of *htt* mutant axons display transient extension of a process into the injury gap but no stable forward growth. (f) Quantification of the time at which forward growth occurs after transection in *htt* siblings (n=10) and *htt* mutants (n=11). Differences measured using Welch’s t-test (***p=0.0002). (g) Quantification of branching events within injury gap of *htt* siblings (n=9) and *htt* mutants (n=11). Branches counted were defined as divergent events visible on axon while axon was in the injury gap. Injury gap was defined as the edge of the distal axon tip. Difference in number of branches determined using Welch’s t-test (*p=0.0412). (h) Quantification of axons that initiate extension within 24 hpt. Extension defined by forward movement for at least three consecutive timelapse frames. Difference in proportions between groups measured with a Fischer’s Exact t-test (*p=.0302). (i) Quantification of growth velocity from 24-48 hpt of *htt* sibling (n=7) and *htt* mutant (n=5) axons, (** p= 0.0011, Welch’s t-test). dpf: days post fertilization; hpt: hours post transection. +/ :sample includes both +/+ and +/- genotypes.

To further explore this possibility, we tracked individual M-cell axonal regrowth trajectories from 24-96 hpt, taking images at 24 h intervals. Analyses of these images demonstrate that in *htt* siblings ^(+/+;-/+)^, M-cell axons regrew to ∼10% pre-injury length by 24 hpf and then increased growth rates between 48-96 hpt to reach ∼80% of their pre-injury length (n=7/7; Figure 3b), indistinguishable from what we observe in wild-type animals (Figure 1b). In contrast, *htt* ^-/-^ axons display several distinctive trajectories that resulted in various deficit in axonal regrowth at 96 hpt. Specifically, we find that 60% (n=6) of *htt* ^-/-^ axons reached between 20-40% pre-injury length at 96 hpt (orange and magenta curves in Figure 3b), while 40% (n=4) of all *htt* mutant axons analyzed in this experiment reached less than 5% of pre-injury length (red curve in Figure 3b). To characterize dynamics of the most severely affected axons (reaching less than 5% of pre-injury length), we performed *in vivo* time lapse imaging between 4-24 hpt. This revealed that the majority of *htt* ^-/-^ regenerating M-cell axons (n=11/18) were motile yet displayed delayed entry into the injury gap (n=11/18). In wild-type siblings, this entry begins at 7.2 ± 2.1 hpt (Figure 3c, f; Supplemental Movie 3), while in *htt* ^-/-^ axons this process is delayed by ∼50% to 11.7 ± 2.4 hpt (Figure 3f). In addition, we find that relative to regenerating M-cell axons in wild-type siblings (1.1 branching events, n=9), a significant subset of *htt* ^-/-^ axons that form a motile growth cone (n=11), display increased and often transient axonal branching while navigating the injury site (4.2 branching events), ultimately reducing regrowth speed and accuracy (Figure 3d, g; Supplemental Movie 4). The most severe phenotype we observed in 38.8% of *htt* mutants (n=7/18) is characterized by repeated, transient M-cell axon extension and retraction but lack initiation of stable forward growth, resulting in a complete failure to enter the injury site (Figure 3e, h; Supplemental Movie 5).

In addition to these defects early during the regeneration process, our analysis suggested a failure of regenerating *htt* ^-/-^ axons to increase the speed after 24 hpt (n=4/10), as well as a delay of increased regeneration rate until after 48 hpt (n=2/10) (Figure 3b). To determine if *htt* is indeed required for this transition into higher speed, long-range growth, we imaged axons between 24-48 hpt, selectively choosing *htt* ^-/-^ axons that successfully navigated the injury site. We find that in comparison to wildtype siblings (n=7, 15.83±3.76 µm/hr), *htt* mutants display slower growth velocity during extension after traversing the injury gap (n=5, 8.194±1.36 µm/hr; Figure 3i; Supplemental Movie 6, 7). Combined, our analyses show that *htt* mutants exhibit distinct phenotypes when axons enter and traverse the injury gap and later when they increase and sustain higher growth rates, strongly supporting the idea that *htt* regulates multiple, distinct steps of spontaneous M-cell axon regeneration.

### *Htt* is dispensable for Peripheral Motor axon and Optic Nerve regeneration

Similar to other vertebrates, *htt* mRNA is detectable throughout the zebrafish nervous system [54]. We therefore asked whether *htt* acts broadly to promote axon regeneration in the central and peripheral nervous system, or whether it plays a more selective role in M-cell spinal cord regeneration. We first examined whether *htt* plays a role in peripheral motor axon regeneration. We previously established a spinal motor nerve regeneration assay (Rosenberg et al., 2012), in which we use laser axotomy to fully transect ventral motor nerves, and measure regeneration at 48 hpt (Figure 4a). For this we use the *tg*:*mnx1*:GFP transgenic line which labels segmentally arranged spinal motor nerves, each consisting of ∼70 individual motor neurons whose axons project to their ventral muscle targets (Figure 4a) [55]. At 5 dpf, spinal motor axons have formed mature neuromuscular junctions with their muscle targets, are myelinated, and are critical to mediate various behaviors including the acoustic startle response and prey catching [56–58]. Prior to nerve transection at 5 dpf, *mnx1*:GFP positive spinal motor nerves in *htt* ^-/-^ animals were indistinguishable from those in wild-type siblings (Supp. Figure 1a-b). We have previously shown that by 48 hours following complete nerve transection, the majority of wildtype nerves regrew multiple fascicles into their former muscular target (Figure 4b) [59]. When compared to *htt* siblings, motor nerves in *htt* mutants display a similar degree of regrowth (Figure 4c), demonstrating that, in zebrafish, *htt* is dispensable for peripheral motor axon regeneration.

**Figure 4:**
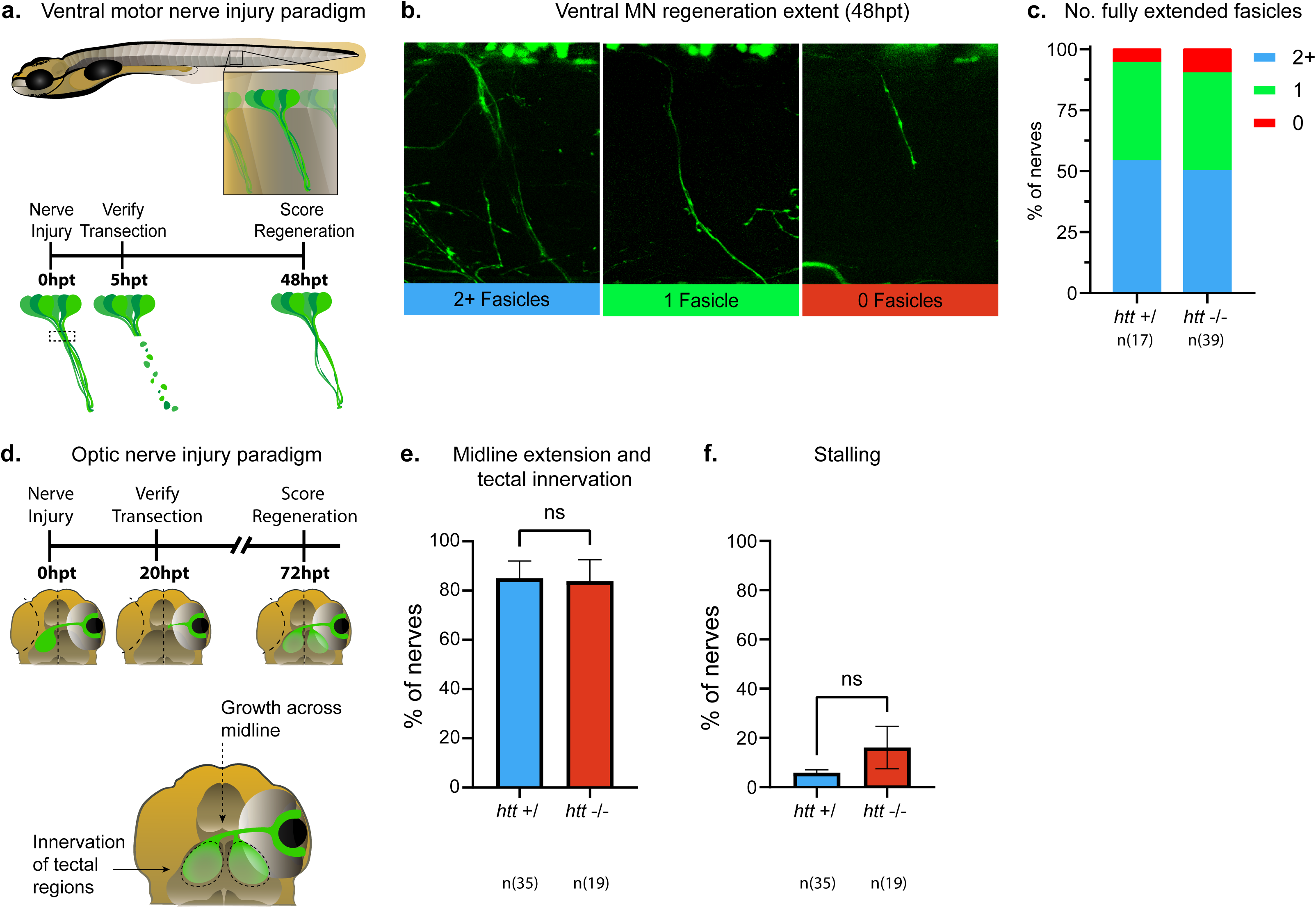
Htt is dispensable for motor nerve and optic nerve regeneration. (a) Schematic of motor nerve injury paradigm. Ventral motor nerves are located within the spinal cord and project axons into peripheral muscle segments along the zebrafish trunk. Nerves are laser-transected, complete transection confirmed at 5 hpt, and regeneration assessed at 48 hpt. (b) Representative images of 0, 1, and 2+ regenerating fascicles at 48 hpt. (c) Quantification of extent of nerve regrowth between *htt* siblings (n=17) and *htt* mutants (n=39). No significant difference observed in regeneration of motor nerves (p=0.9123, Fischer’s Exact test). (d) Schematic of optic nerve injury paradigm. Optic nerves are mechanically transected, complete transection confirmed at 20-24 hpt, and regeneration assessed at 72 hpt. Robust regeneration is a combined score of nerves that display extension across the midline of the brain and reinnervation of tectal regions. (e) Quantification of nerves that cross the midline and reinnervate the tecta at 72 hpt in *htt* siblings (n=35) and *htt* mutants (n=19). No significant difference in proportions between groups measured (p=0.5000, paired t-test). (f) Quantification of nerves displaying stalled axons in htt siblings and htt mutants. No significant difference in proportions between groups (p=0.3103, paired t-test). +/ :sample includes both +/+ and +/- genotypes.

We next asked whether *htt* promotes spontaneous axon regeneration throughout the CNS or selectively in M-cells. Following optic nerve crush in mice, axon regeneration is suppressed in HTT/PTEN-KO, regeneration-competent Retinal Ganglion Cells (RGCs) [49], however due to the poor regenerative capacity in mammals, the role of *huntingtin* in spontaneous i.e. unforced optic nerve regeneration is currently unknown. We therefore asked whether zebrafish *htt* is required for spontaneous optic nerve regeneration. We have previously established a larval assay to measure optic nerve regeneration following complete optic nerve transection [60]. For this we used *tg:Isl2b-GFP* to label RGC soma and axons [61], and performed optic nerve transections at 5 dpf (Figure 4d). To assess *htt*’s role in optic nerve regeneration, we quantified regeneration in *htt* ^+/+; +/-^ siblings and *htt* ^-/-^. Similar to wild-type siblings at 72 hpt, around 80% of transected *htt* ^-/-^ optic nerves had crossed the CNS midline and reinnervated the optic tecta without displaying significant levels of premature stalling (Figure 4e, f). Thus, *htt* is dispensable for optic nerve regeneration, yet plays a critical role in axon regeneration selectively for spinal cord M-cell axons.

### *Huntingtin* modulates initiation of the transcriptional injury response

Our data reveals that *htt* promotes early steps of spinal M-cell axon regeneration enabling regenerating axons to become motile and traverse the injury site (Figure 3). We therefore wondered whether *htt* functions to activate the transcriptional injury response, key for neurons to respond to axonal damage by initiating growth-promoting gene programs [62, 63]. To test this idea we performed *in situ* hybridization at 24 hpt to measured M-cell specific mRNA upregulation of two cross-species validated <u>R</u>egeneration <u>A</u>ssociated <u>G</u>enes (RAGs), *atf3* and *gap-43* [64] (Figure 5a). As expected, in non-transected M-cell soma of wildtype siblings we did not detect significant upregulation of *atf3* and *gap-43* mRNA levels (Figure 5b). Conversely, in 100% of transected wildtype M-cells analyzed we find significantly elevated *atf3* mRNA levels (Mean Intensity ΔF compared to non-transected internal control: 4015 ± 1346 a.u., n=14; Figure 5b). In contrast, in transected *htt* ^-/-^ M-cells, we failed to detect significant upregulation of *atf3* expression (Mean Intensity ΔF transected versus non transected: 391.9 ± 504.4 a.u., n=14; Figure 5B-D). Similarly, *gap-43* expression is robustly upregulated in transected wildtype M-cell somas at 24hpt (Mean Intensity ΔF 3058 ± 695.9 a.u., n=10), while in *htt* mutants, *gap-43* upregulation is greatly blunted (Figure 5F-H; Mean Intensity ΔF-21.73 ± 274.1 a.u.; n=8). In contrast to *htt* mutants, *celsr3* mutants cross the injury gap at wild-type like speed and only thereafter fail to increase regeneration speed. We detect robust M-cell *atf3* mRNA upregulation in *celsr3* mutants, supporting the notion that *celsr3* is dispensable during the very first regeneration steps and acts selectively after injury site crossing (Supplemental Figure 2 a-c). Thus, in wildtype M-cells, injury induced upregulation of *atf3* or *gap-43* mRNA correlates with regenerating growth cones traversing the injury site. Meanwhile, *htt* ^-/-^ M-cells fail to upregulate atf*3* or *gap-43* mRNA along with the failure of their axons to traverse the injury gap. Combined, our results demonstrate that *htt* promotes early steps of spinal cord regeneration and provide compelling evidence that this occurs at least in part by initiating the transcriptional injury response.

**Figure 5:**
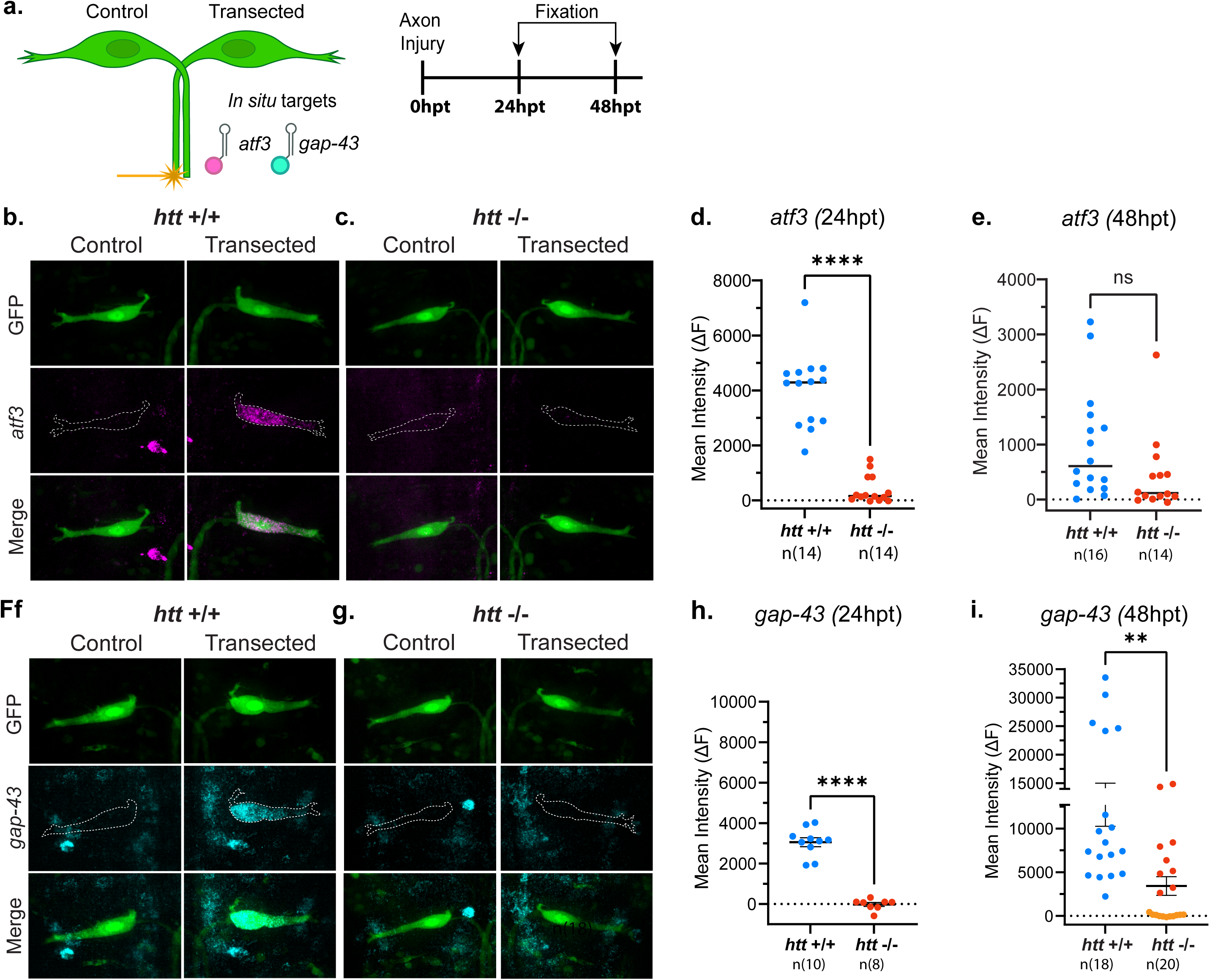
*Huntingtin* modulates prompt initiation of the transcriptional injury response in M-cells. (a) Schematic of *in situ* hybridization assay on the M-cell with gene targets *atf3* (magenta) and *gap-43* (cyan). (b-c) Images of *atf3* upregulation within injured M-cell (24hpt) with uninjured (control) M-cell in *htt* +/+ and *htt* -/- examples. (d) Quantification of difference in *atf3* Mean Fluorescence Intensity between injured and uninjured M-cells of *htt* siblings (n=14) and *htt* mutants (n=14) at 24 hpt, (**** p=>0.0001, Welch’s t-test). (e) Quantification of difference in *atf3* Mean Fluorescence Intensity between injured and uninjured M-cells of *htt* siblings (n=16) and *htt* mutants (n=14) at 48 hpt, (p=0.0873, Welch’s t-test). (F-G) Images of *gap-43* upregulation within injured M-cell (24 hpt) with uninjured M-cell in *htt* +/+ and *htt* -/- examples. (h) Quantification of difference in *gap-43* Mean Fluorescence Intensity between injured and uninjured M-cells of *htt* siblings (n=10) and *htt* mutants (n=8) at 24 hpt, (**** p=>0.0001, Welch’s t-test). (i) Quantification of difference in *gap-43* Mean Fluorescence Intensity between injured and uninjured M-cells of *htt* siblings (n=20) and *htt* mutants (n=18) at 48 hpt, (* p=0.0412, Welch’s t-test).*htt* mutants displaying little *gap-43* expression at 48hpt colored orange; *htt* mutants displaying higher *gap-43* expression at 48hpt colored red.

Our static and dynamic analyses of M-cell axon regeneration in *htt* mutants revealed that a significant portion of M-cell axons, while unable to reach pre-injury length, regrew significantly past the injury gap (27.6%, n=8), including axons which grew up to 40% pre-injury length (Figures 2, 3). While we observe a significant reduction in *atf3* and *gap-43* mRNA upregulation in nearly 100% of *htt* mutants at 24 hpt, it is possible that *atf3* and *gap-43* mRNA upregulation, rather than being abolished, is delayed. While this delay likely accounts for the inability of transected axons to reach their full pre-injury regrowth levels, it might be sufficient for regenerating M-cell to initiate regrowth later and extend significant distances beyond the injury gap. To test this, we examined *atf3* and *gap-43* mRNA in M-cells at 48 rather than 24 hpt. In wild type animals, *atf3* mRNA expression remains upregulated in transected M-cells and *gap-43* mRNA expression was markedly higher at 48 hpt relative to levels at 24 hpt (compare Figure 5d to e; h to i). Furthermore, when compared to transected wild type M-cells, *atf3* expression levels in *htt* ^-/-^ M-cells were indistinguishable compared to those in wild type siblings (Figure 5e), while levels of *gap-43* mRNA expression bifurcated into a subset of mutant M-cells that still displayed no significant expression (orange data points in Figure 5i), and into a subset that now displayed expression level comparable to those in wildtype siblings (red data points in Figure 5i). We wondered whether this bifurcation in *gap-43* expression was correlated with *htt* ^-/-^ axon regrowth extent. To address this, we examined a subset of *htt* -/- and *htt +/+* samples and correlated *gap-43* expression with axon regrowth length at 48 hpt (Supplemental Figure 3) We find that *htt* mutants with little detectable expression of *gap-43* also displayed no regrowth (n=2, orange dots) while *htt* mutants with *gap-43* expression comparable to *htt* +/+ siblings did extend beyond the site of injury (n=4, red dots). Next, we sought to determine whether comparable *gap-43* mRNA expression in *htt* mutants at 48hpt is sufficient for wildtype-like regeneration length. We find that *htt* mutants expressing wildtype-like levels of *gap-43* at 48 hpt still exhibit shorter regenerative axonal length (Supplemental Figure 3), suggesting that additional factors such as the expression onset time or duration play critical roles. Combined our data provide compelling evidence for a mechanism by which *htt,* rather than being essential to initiate a transcriptional injury response, ensures its timely onset, thereby promoting a greater extent of long-range axonal regeneration.

### *Htt* regulates M-cell regeneration in a distance-dependent manner

One prediction from this model is that accelerating the onset of the transcriptional injury response should improve M-cell axon regeneration in *htt* mutants. The transcriptional injury response depends on dynein/microtubule mediated retrograde transport of injury signals from the site of axotomy towards the soma [65]. Combined with the well documented role of HTT in bi-directional axonal transport, we decided to test this prediction by reducing the distance between injury site and soma, thereby accelerating the onset of RAG upregulation. To reduce injury site to soma distance, rather than transecting M-cell axon approximately 780 µm distal to the cell body as done before (distal location), we transected wildtype M-cell axons at a more proximal location, at ∼280 µm from the soma (Figure 6a; proximal location). Comparing *atf3* expression at 12hpt in M-cells transected at the distal location versus the proximal location revealed significant *atf3* mRNA upregulation in proximal transected M-cells versus those transected at the distal location, confirming that reducing injury site to soma distance accelerates transcriptional injury response onset (Figure 6B). We then tested whether reducing injury site to soma distance in *htt* mutants leads to increased *atf3* expression levels. Consistent with our previous results (Figure 5), in response to a distal site transection, the majority of *htt ^-/-^* M-cells fail to display appreciable *atf3* expression at 24 hpt. In contrast, following proximal site transection in *htt* mutants induced *atf3* expression to levels comparable to wildtype siblings (Figure 6c). Lastly, we asked whether accelerated onset of the transcriptional injury response improves M-cell axon regeneration in *htt* mutants. In fact, when we compared axonal length following distal or proximal transections, we found that this did not alter regeneration in wildtype siblings, but significantly improved regeneration in *htt* mutants, resulting in a mean fourfold increase from 15% to over 60% of pre-injury length (Figure 6d).

**Figure 6:**
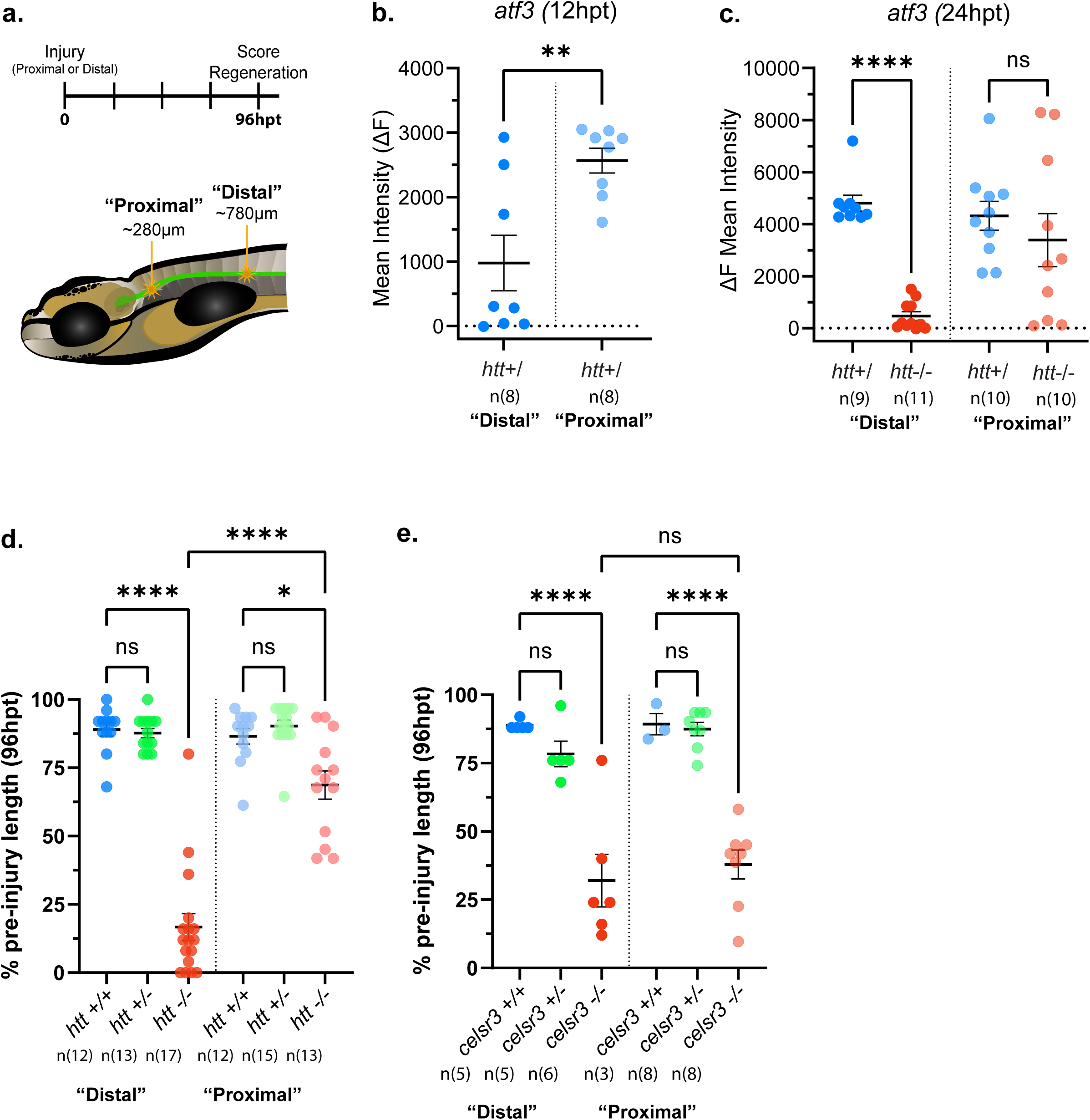
*Htt* regulates M-cell regeneration in a distance-dependent manner. (a) Schematic of M-cell transection at proximal (approx. 280 µm from cell soma) and distal (approx. 780 µm from cell soma) locations. (b) Quantification of *atf3* expression at 12 hpt of wildtype M-cells with axons transected at the proximal (n=8) or distal (n=8) locations. At 12 hpt, *atf3* expression is significantly lower in M-cells transected at the distal injury location compared to those transected at the proximal injury location (p=0.0075, Welch’s t-test). (c) Quantification of *atf3* expression at 24 hpt of M-cells with axons transected at the proximal or distal locations of *htt* +/ and *httv*-/-. *Htt* -/- transected distally display low Δ Mean Fluorescence compared to *htt* +/ (****p<0.0001, Sidak’s). However, *htt* -/- transected at the proximal location display no significant difference in Δ Mean Fluorescence compared to *htt* +/ (p=0.7348, Sidak’s multiple comparisons). (d) Quantification of axon regrowth (96hpt) of *htt* +/+, *htt* +/- and *htt* -/- at proximal and distal locations. Transections at the distal location result in regeneration deficits in *htt* -/- compared to *htt* +/+ controls (***p< 0.0001, Tukey’s multiple comparisons). Transections at the proximal location resulted in less severe deficits in *htt* -/- compared to *htt* +/+ controls (**p=0.0061, Tukey’s multiple comparisons) and a significant increase in regeneration compared to distally transected *htt* -/- (****p< 0.0001, Tukey’s multiple comparisons). (e) Quantification of axon regrowth (96 hpt) of *celsr3* +/+, *celsr3* +/-, and *celsr3* -/- at proximal and distal locations. Transections at the distal location result in regeneration deficits in *celsr3* -/- compared to *celsr3* +/+ controls (****p< 0.0001, Tukey’s multiple comparisons). Transections at the proximal location still resulted in deficits in *celsr3* -/- compared to *celsr3* +/+ controls (***p=0.0004, Tukey’s multiple comparisons) and no significant change in extent of regeneration compared to distally transected *celsr3* -/- (p= 0.4331, Tukey’s multiple comparisons).

A second prediction from this model is that axon regeneration promoted by genes that act at a later step in the regeneration process, in particular after the onset of the transcriptional injury response, would be unaffected by reducing injury site to soma distance. To test this prediction, we assayed the effects of proximal M-cell transections in *celsr3* mutants. After onset of the transcriptional injury response *celsr3^-/-^* M-cell axons enter and traverse the injury gap at speeds indistinguishable from wild-type axons, and only after crossing the injury gap fail to increase growth rates between 24 and 48 hpt (Fig. 2f, h), consistent with the idea that *celsr3* functions after the onset of the transcriptional injury response. In fact, we find that unlike *htt* mutants, proximal transection of *celsr3^-/-^* M-cell axons fails to increase axonal regeneration (Figure 6e). Combined these results provide compelling support for a mechanism by which *htt* functions early during M-cell axon regeneration to accelerate the onset of the transcriptional injury response critical to ensure high efficiency, long-range axon regeneration.

### Knockdown of *htt* interacting axonal transport proteins recapitulates *htt* regeneration defects

The ability to partially restore axonal regeneration in *htt* mutants by reducing axotomy-to-soma distance suggests an *htt*-dependent transport role in promoting M-cell regeneration. Acting via specialized protein binding partners, *htt* has been implicated in a wide range of cellular functions, including transcriptional regulation [48, 49] such as through *tp53* [39, 40], as well as autophagy, endocytosis and bidirectional axonal transport [38, 47]. To determine whether *htt* indeed mediates long-range spinal cord M-cell regeneration through axonal transport, we tested a diverse collection of *htt* interactors for potential roles in long-range axon regeneration. Given the large number of over 300 interacting proteins, we examined a representative subset of *htt* interactors (Figure 7a) involved in various cellular functions including gene regulation, trafficking/transport and endocytosis. To test for functional *in vivo* roles of interactors we used available genetic mutants as well as a well established F0 CRISPR/Cas9 genetic knock down strategy (Figure 7b; for details on the CRISPR/Cas9 strategy, see Materials and Methods). We find that genetic mutants of the transcription factor *tp53* (Figure 7c), the *hip14* palmitoyl transferase (Figure 7d), and the selective autophagy receptor *Optineurin* (optn) (Figure 7e) when assayed for M-cell axon regeneration, did not display significant long-range regeneration defects. Knockdown of the multifunctional huntingtin-interacting protein 1 (Hip1) resulted in a mild yet significant reduction in axonal regrowth without causing a severe reduction in regrowth length (around 20% pre-injury length characteristic for *htt* mutants) (Figure 7f). In contrast, knock down of genes selectively involved in vesicle sorting and motor-based axon transport displayed significant deficits in M-Cell axon regeneration. Specifically, knock down of huntingtin associated protein 40 (hap*40*), which complexes with HTT to mediate sorting and trafficking of rab5+ vesicles [45, 46], adaptor protein huntingtin associated protein 1 (*hap1) and* protein arginine methyltransferase 6 (*Prmt6*), both of which regulate the speed and direction of motor-based axon transport ([42–44, 66], resulted in significant deficits in axon regeneration (Figure 7g-i). Thus, these results provide genetic *in vivo* support for a model by which *htt* regulates the onset of the transcriptional injury response via a mechanism of axonal vesicle transport.

**Figure 7:**
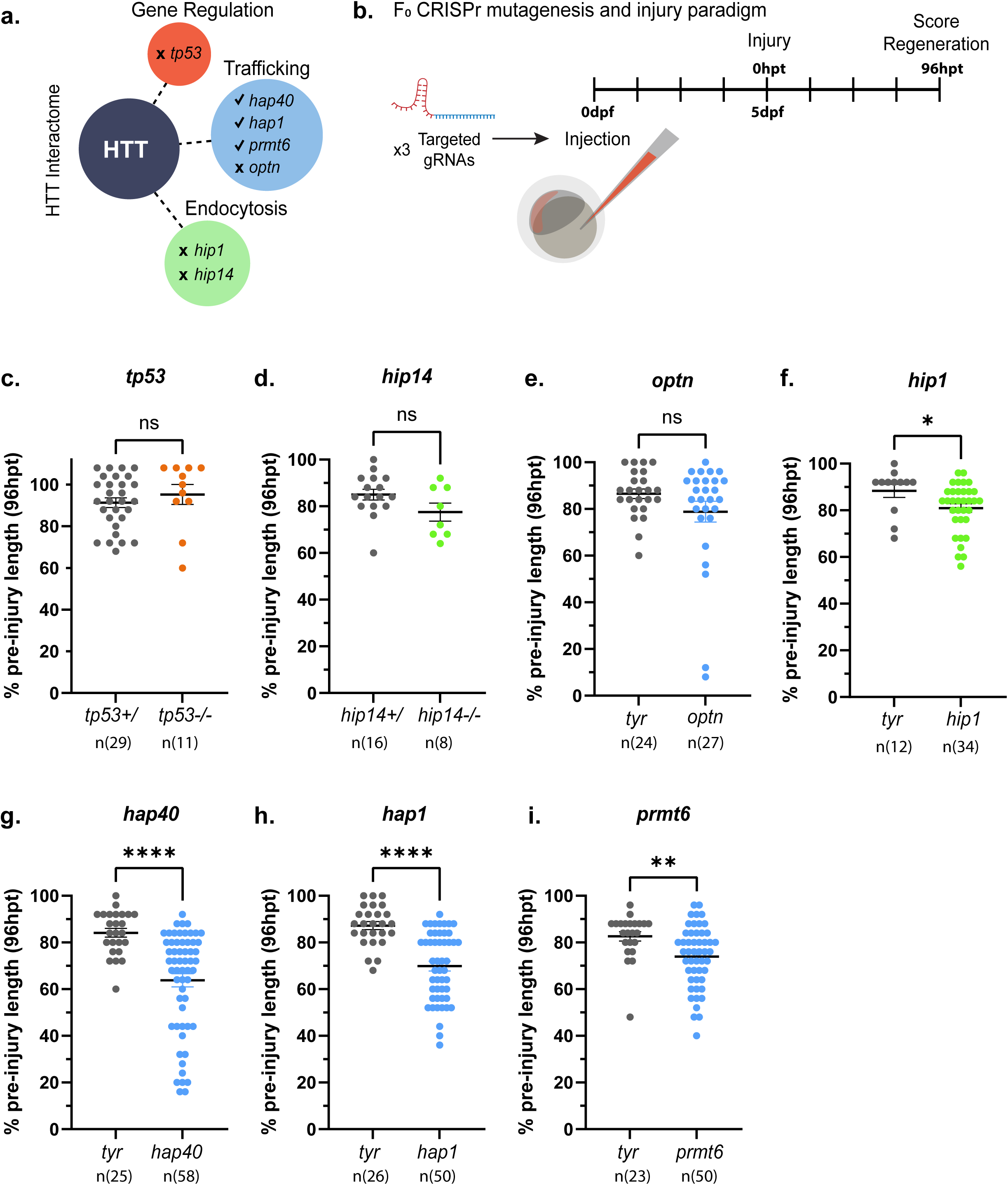
HTT-interacting proteins involved in axonal transport also display deficits in M Cell regeneration. (a) Schematic of HTT interactome including large functional classes of tested interacting proteins. (b) Schematic of F0 CRISPr-Cas9 gene editing paradigm. Target gRNA sequences were created against selected genes of interest, injected into 1-cell wildtype embryos and larvae were raised to 5 dpf. Axons were transected and subsequent scoring of regeneration occurred at 96 hpt. (c) Quantification of axon regeneration at 96 hpt in *tp53* siblings (n=29) and *tp53* mutants (n=17) (p=0.4712, Welch’s t-test). (d) Quantification of axon regeneration at 96 hpt in *hip14* siblings (n=16) and *hip14* mutants (n=8) (p=0.1206, Welch’s t-test). (e-i) Quantification of regeneration length of larvae injected with guides targeting (e) *optn* (p=0.1250 Welch’s t-test, n=27), (f) *hip1* (*p=0.0381 Welch’s t-test, n=34), (g) *hap40* (****p<0.0001, Welch’s t-test n=58), (h) *hap1*(****p<0.0001 Welch’s t-test, n=50), and (i) *prmt6* (**p=0.0026 Welch’s t-test, n=50). Controls consisted of wildtype larvae injected with gRNA targeting the *tyrosinase* (*tyr*) gene, which has no effect on M-cell regeneration.

## Discussion

Unaided or spontaneous CNS axon regeneration, long thought to only occur in non-mammalian vertebrates, has by now been documented in adults of a growing number of mammalian species [17–19]. This argues that in parallel to well established inhibitory pathways, evolutionary conserved genetic programs promoting robust, functional CNS regeneration exist. Yet despite their relevance, the complement of pro-regenerative factors and the mechanisms by which they overcome inhibition, enable regenerating CNS axons to extend along their preinjury path, and reinnervate their original synaptic targets is extremely limited. Here we combine the robust CNS regeneration capacity of zebrafish with its optical transparency and a quantifiable, single axon resolution assay for spinal cord regeneration. We first detail the stepwise and dynamic morphological changes occurring at the regenerating axon tip following complete transection *in vivo*, including rebuilding a motile growth cone, probing and traversing the injury site, and then initiating long-range regrowth. Using this foundation, we determine roles for two pro-regenerative genes isolated from a screen, *htt* and *celsr3,* at overlapping and different steps of the regenerative process. We show that the speed of axonal regeneration is not uniform along the regenerative route but rather tunable, and that *celsr3* is a critical component of a switch enabling axons to precipitously increase speed once they completed traversing the injury site. Moreover, we demonstrate that *htt* plays multiple roles in axon regeneration, including ensuring a timely transcriptional response consistent with a *htt*-dependent critical window of regenerative competency. Finally, we provide compelling functional evidence that *htt* promotes long-range spinal cord regeneration through axonal transport. Taken together, we provide a first single axon resolution, minute by minute, timeline describing the dynamic behaviors of regenerating vertebrate spinal cord axons in their native environment, and place two pro-regenerative genes with selective and partially overlapping functions into a genetic program underlying spontaneous spinal cord regeneration.

### Multiple genetic factors promote axon motility at different times during axon regeneration

Our experiments discern divergent temporal requirements of *celsr3* and *htt* following injury despite both mutants exhibiting similar axon regeneration deficits at 96hpt. *Htt* is already required at the onset of axon reorganization, as mutants display both failure and delay in the formation of a motile process that in wildtype animals extends around 7hpt. The timing of this reorganization likely precedes significant responses from the cell body, suggesting *htt* acts locally to regulate these early events. The dynamics of prolonged extension and retraction resembled *in vitro* characterizations of dystrophic growth cones [21, 67], which occur when axons fail to properly reorganize the cytoskeleton. HTT is known to directly bind to tubulin subunits as well as Tau and Dynein, both known to promote microtubule stability [68, 69]. Future studies to assess the internal structure of the proximal axon in *htt* mutants, including through electron microscopy, may reveal additional roles for *htt* underlying the structural organization and formation of nascent regenerative growth cones.

Compared to the pathways underlying growth cone formation and early sprouting, much less is understood about the mechanisms promoting sustained, efficient long-range axon elongation. Our results reveal a selective role for *celsr3* in later, fast axon extension, demonstrating that the molecular makeup of the regenerative growth cone evolves throughout the course of regeneration and that the increase in axon speed is essential for long-range growth. What causes this switch in extension velocity? One hypothesis is that the transcriptional response to injury produces additional components targeted to the regenerating growth cone initially not available there, increasing efficiency. Supporting this, the timing of our increased extension speed is concomitant with cell body upregulation of several regeneration-aiding genes, such as *gap-43*. GAP-43 and Celsr3 itself are located in growth cones during development [35, 70]. *Htt* mutants secondarily fail to achieve increased speed of extension, though whether this reflects a shared or parallel mechanism with *celsr3* is unknown. Notably, the M-cell axon extends at speeds of 100 µm/hr during development [71], suggesting that regenerative axon growth is much less efficient, even in wildtype animals. Whether this difference is due to further missing factors within the regenerative growth cone or factors in post-developmental environmental substrates remains unresolved. Regardless, we show that control of regenerative efficiency has marked effects on the degree of regeneration, which agrees with key differences observed between CNS and PNS axon extension in rodents [72].

### A competency window for long-range axon regeneration

Cellular responses to injury occur over several timescales, where early signals, such as calcium signaling from retrograde propagation reach the cell body within minutes after injury, while signals dependent on transport by motor proteins likely take hours to arrive and induce gene expression [62]. That a timely onset of the transcription injury response might be critical for successful long-range axonal regeneration has been suggested [73], yet has not been tested using defined genetic mutants. Our results using *htt* mutants reveal that a significant subset of regenerating M-cells express *aft3* and *gap-43* mRNA levels comparable to wild type siblings at 48hpt but not at 24hpt, when they linger at baseline levels (Figure 5). Importantly, expression levels at 24 hpt correlate with achieving long-range regeneration (Figure 1h) in wild type siblings, while high expression levels at 48 hpt fail to achieve comparable long-range regeneration in *htt* mutants. This reveals that robust, yet delayed upregulation of injury response genes is insufficient for axons to fully re-extend, consistent with the idea that injury induced molecular events operate in a critical temporal window. Moreover, our results support the idea that injury response latency is a critical predictor specifically for long-range spinal cord regeneration.

### HTT functions as a distance-dependent regulator of axon regeneration

Our results demonstrate that while *htt* is fully dispensable for regeneration of shorter PNS and CNS axons (Figure 4), and partially dispensable even for M-cell regeneration following transection at ∼280 µm (Figure 6), *htt* is critically required for long range regeneration when transection occurs at ∼780 µm from the cell soma. In strong support of a distance-dependent rather than cell type specific regulator, we find that knockdown of HTT-interacting proteins such as HAP40 and HAP1 which play defined roles in promoting retrograding vesicular trafficking through endosome sorting and modulation of molecular motors recapitulate the *htt* mutant M-cell regeneration phenotype. While our experiments do not exclude all mechanisms of gene regulation to be critical for M-cell regeneration, our findings support a primary role of HTT regulating axonal trafficking. This is consistent with a model by which *htt*-dependent axonal trafficking controls the onset of the neuronal injury response, likely around the time when injury signal proteins arrive reach the soma. The distance between the cell soma and site of injury is known to be an important determinant of regenerative capacity across several neural populations, in particular for supraspinal neurons whose axons extend over very long distances [74]. Thus, while the phenomenon of distance-dependent regenerative capacity is well-established, our work identifies *htt* as a previously unappreciated determinant of distance-dependent regeneration. Currently, models predict that neurons detect and encode distances on the dynamics of motor-based signals. One model uses a time-delay between early priming signals and subsequently trafficked signals, while a second model predicts an oscillatory delay between anterograde and retrograde trafficking events [63, 75, 76]. Our results demonstrating a time dependent role for *htt* are consistent with *htt* playing a role in either of these proposed mechanisms, and additional experiments are required to fully define the mechanism by which HTT links axonal trafficking to regenerative capacity. The current view is that HTT might be involved in multiple mechanisms [38, 48, 49] promoting axon regeneration. Thus, our work uncovers a distinct and previously unappreciated perspective on the role of endogenous HTT regulating axonal trafficking through mechanism dependent on soma to injury site distance rather than a cell type specificity.

## Acknowledgments

We would like to thank Dr. Andrea Stout of the UPenn Cell and Developmental Biology Microscopy Core. We also thank all members of the Granato laboratory for comments and discussion.

## Methods

### Ethics Statement: (IACUC protocol)

Protocols and procedures involving zebrafish (*Danio rerio*) husbandry and care are in accordance with the University of Pennsylvania Institutional Animal Care and Use Committee regulations. Veterinary care is under the supervision of the University Laboratory Animal Resources (ULAR) of the University of Pennsylvania.

### Zebrafish strains and animal care

Transgenic lines were maintained in TU or TLF backgrounds and raised as described in [77]. All embryos were collected a few hours after fertilization and raised at either 28°C or 29°C. At the time of experiments, zebrafish have not yet determined sex. To visualize the Mauthner cell, tg (Tol-056-GFP) was used, as described in [33]. tg(isl2b-GFP) was used to visualize Retinal ganglion cells (RGCs) [61]. To visualize motor neurons, tg (hb9-GFP) was used [78].

The mutant strains used are as follows: *celsr3 ^fh339^*[22], *tp53^M214K^* [79], *hip14^p174^* [80], and htt*^p423^.* Genotyping for *tp53, hip14, celsr3,* and *htt* alleles was performed using KASP primers with proprietary primer sequences (LGC Genomics) designed against the sequences listed in (Table 2).

**Table 1:**
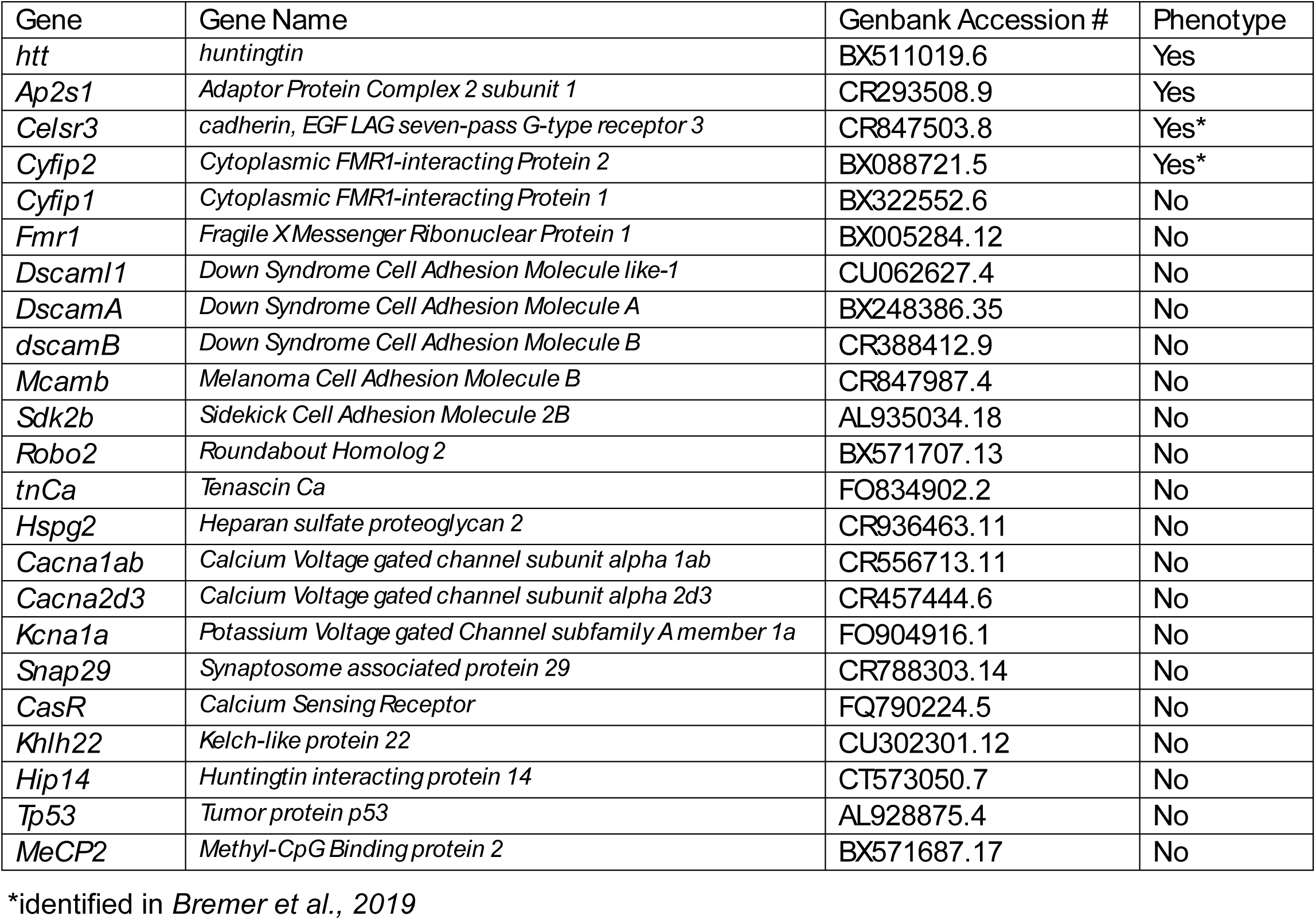
Screen for genetic factors for long-distance M-cell regeneration.

**Table 2:**
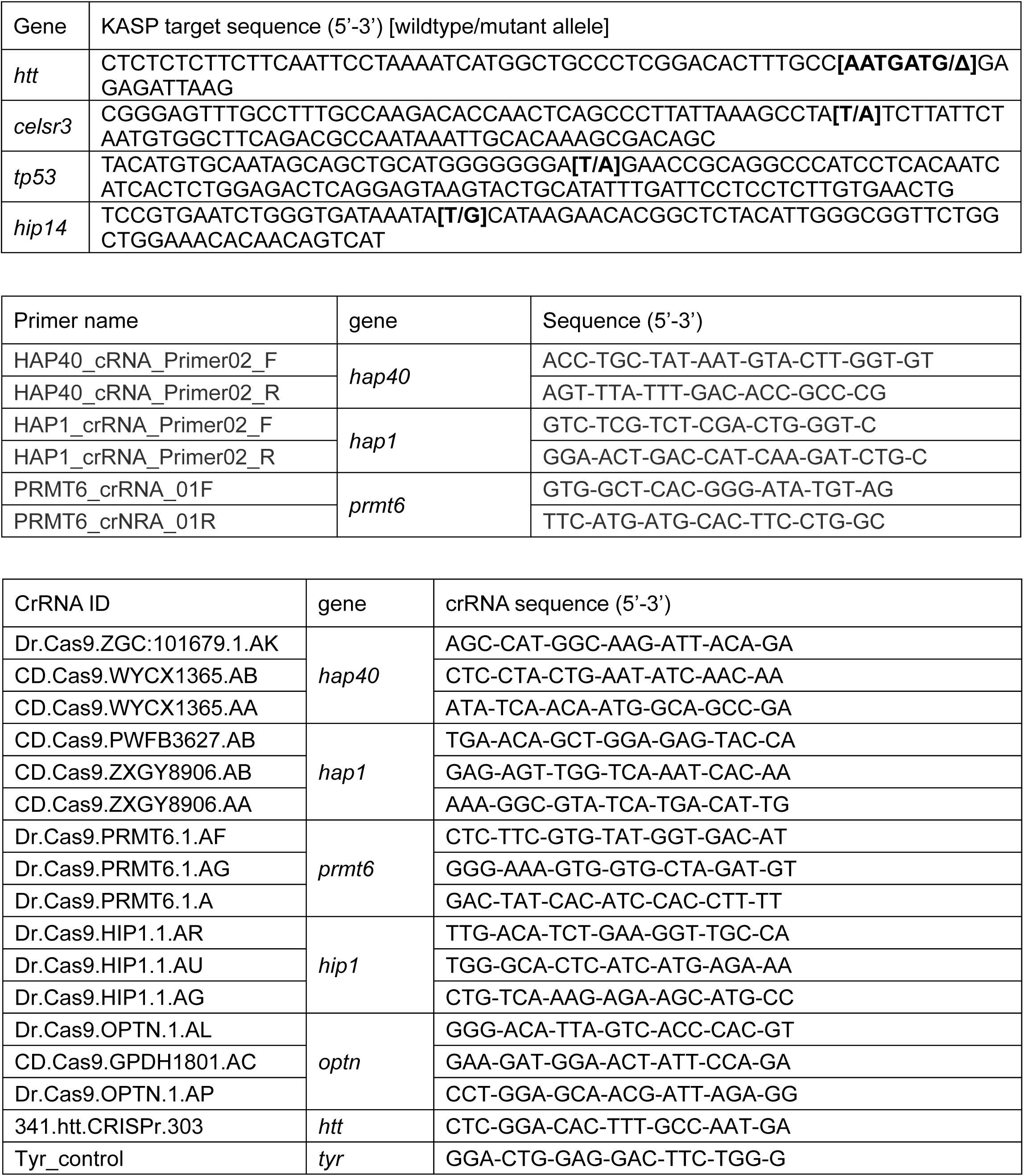
Primer and CRISPr guide sequences used in this study.

### Generation of htt*^p423^*mutant allele

*Htt pa423* allele generated through CRISPr-cas9 mutagenesis. A crRNA guide, sequence 5’-GGA-CTG-GAG-GAC-TTC-TGG-G-3’ targeted 565bp-571bp in exon 6 of the *zhtt* locus, creating a premature stop at 195aa. CrRNA was complexed with tracrRNA (IDT, #1072533) to create an sgRNA guide then complexed with Cas9 protein (IDT, #1081061) and injected into 1-cell stage embryos. F_0_ injected larvae were raised and outcrossed to establish heterozygous F_1_ carrier lines. Heterozygous F_1_ carrier alleles were identified through PCR genotyping and sequencing.

### M-cell Transection Assay

At 5dpf, larvae were anesthetized in 0.002% Tricaine E3 and mounted on their side in 1.5% agarose/E3 on a coverslip-bottom dish (MatTek). M-cell axon transections were performed on a VT-iSM camera confocal with Metamorph Imaging Software. Repeated bouts (5-7 times) of 330msec 355nm iLAS laser exposure was applied to a region approximately 3×3µm over one Mauthner axon at the location of the 9^th^ trunk hemisegment. Full transection and appropriate retraction of the axon was assessed 20-30min after initial laser exposure. Only larvae with complete separation of the distal and proximal tip and without retraction beyond the boundary of the 9^th^ somite were used to assess regeneration. For proximal transections, the same injury conditions and verification was applied at the 3^rd^ somite. Following injury, larvae were dismounted from agarose and recovered in E3 medium at 28 C until fixation or subsequent imaging at various timepoints. To quantify the extent of regrowth of the transection M Cell axon, larvae were anesthetized in Tricaine and remounted in 1.5% agarose. Imaging and quantification of regrowth was performed on a Spinning Disk Confocal and Slidebook 3.0 software. The site of injury was verified and length was quantified by counting the number of somites from the site of injury that the axon enters and dividing that number by 25 (the average somite length of the M Cell pre-transection).

### Optic Nerve Transection Assay

Optic Nerve transections were performed as described in [60]. To visualize optic nerve regrowth, larvae were raised in 0.2mM phenylthiourea (PTU) E3 medium, covered, to inhibit pigmentation. Larvae were kept at 29°C to reduce slight delays in development from the PTU treatment. At 3dpf, larvae were anesthetized in 0.002% Tricaine PTU/E3, one eye was removed using forceps, and fish were left in Ringer’s solution for 1 hour before being returned to PTU/E3 medium. At 5dpf larvae were again anesthetized in 0.002% Tricaine PTU/E3 and mounted ventral side up in 2.5% agarose on a microscope slide. Optic nerves were visualized under a fluorescent microscope (Olympus SZX16) and transected using a sharpened tungsten needle. Transections were directed at the nerve just after exiting the eye but before the optic chiasm. Larvae were removed from the agarose and again left to recovery for 1 hour in Ringer’s solution before being transferred to PTU/E3. Complete transections were verified between 20-24hpt and larvae with visible tecta labeling or axons extending beyond the chiasm were excluded. Regeneration was scored by mounting larvae in 2.5% agarose and visualized under a fluorescent microscope at 72hpt.

### Ventral Motor Nerve transection assay

At 5dpf, larvae were anesthetized in 0.002% Tricaine E3 and mounted on their side in 1.5% agarose/E3 on a coverslip-bottom dish. Before and after photos of selected ventral motor nerves (VMNs) on a spinning disk microscope (Olympus). Transections were performed on a VT-iSM camera confocal using a 355nm iLAS laser. Repeated bouts (4-6 times) of 330msec laser exposure was applied to a region approximately 3umx10um (width of nerve bundle). 2-3 non-consecutive ventral nerves per larvae were transected. Complete transection was verified at 5hpt and samples with visible uninjured axons or substantial damage to the myotome were excluded. Z-stacks of regenerated nerves were taken at 48hpt and exported to FIJI for visualization and quantification. Degree of regeneration was scored by counting the number of fully extended axon bundles or fascicles and categorized into three groups corresponding to (1) no regrowth/incomplete axon extension, (2) 1 fully extended fascicle, and (3) 2 or more extended fascicles.

### Time-lapse imaging and analysis

Larvae anesthetized in 0.002% tricaine were mounted in 1.2% agarose/E3 in coverslip-bottom dishes (MatTek) filled with E3 medium. Live imaging was done on an Olympus IX81 microscope equipped with a Yokogawa CSU 10 scan head and a Hamamatsu EMCCD camera (model C9100-13) with Slidebook 5.0 Imaging software. The microscope stage was fitted with a LiveCell incubator (Pathology Devices Inc.) set to 28 °C for the duration of imaging. Images acquired at 40x water UAPO/340 immersion objective, 1.15 NA. with Immersol instead of water applied to objective for imaging clarity over time. GFP capture detected with 488nm diode laser. For all timelapses, z-stacks were captured at 20min intervals for 20-24hours. Z-stacks were compressed into maximum projections and exported as 16-bit TIFF images for analysis in FIJI.

Timelapses were background corrected for visual contrast and adjusted for 3D drift. Forward extension was defined as the timepoint after three consecutive frames of axon elongation without retraction. Branching was defined as a clear separation of at least two processes that persist for at least 2 frames. Extension velocity was calculated by measuring displacement of the axon tip using the line tool and dividing the distance with the frame count. If more than one axon tip was present, the fastest growing process was measured.

### Static imaging and analysis

Static images were acquired using an Olympus IX81 microscope with the same parameters as timelapse images. Images acquired spanned about 148µm of M-cell axon length, which included the boundaries of one somite segment, with some overlap between images. Z-stacks were compressed into maximum projections and exported as 16-bit TIFF images for analysis in FIJI. Measurements were taken using the line tool to measure the length of consecutive somite boundaries. Rate increase (Figures 1h and 2g) was determined by measuring the difference in axon length between images taken at 24 and 48hpt within the same animal and divided by 24h. Example images adjusted for brightness and contrast and color assigned in FIJI.

### Fluorescent *in situ* hybridization and analysis

Expression of *atf3* and *gap-43* in larvae was detected using the fluorescent *in situ* hybridization chain reaction (HCR) (Molecular Instruments) and following protocols published in [81]. To summarize, Larvae were anesthetized at relevant timepoints in tricaine and fixed in 4% PFA in 1x PBS and 0.1% TritonX-100 overnight at 4°C. Larvae were washed in 1x PBS+ 0.1% TritonX solution before permeabilization with 100% MeOH at -20°C. 2.0 pmol of predesigned primary probe was added to 500ul of hybridization buffer (Molecular Instruments) and incubated at 37°C overnight. After washing with wash buffer (Molecular Instruments) and 5x SCCT, secondary amplifiers (30pmol each) of 546nm fluorophores were added with 500ul Amplification Buffer (Molecular Instruments). Amplifiers were washed using 5x SCCT and larvae brains were dissected using forceps and mounted in vectashield (Vector Laboratories) overnight. Imaging done on an Olympus spinning disk confocal (described above) with GFP capture and 546nm secondary amplifier visualized with 488nm and 561nm diode lasers respectively. Images were captured as Z-stacks and exported as 16-bit TIFF images for analysis in FIJI.

Z-stacks were compressed into maximum projections. ROI of 111.6 pixel area used to sample Mean Intensity for injured and uninjured M-cell soma in the 561nm channel corresponding to secondary amplifier expression. Mean Intensity Δ was calculated by subtracting the injured value from the uninjured value. Example images adjusted for brightness and contrast and color assigned in FIJI.

### F0-Crispr Injections

Crispr crRNA guides for *hap40, hap1, prmt6, hip1,* and *optn* were designed using the Custom Alt-R CRISPR-Cas9 guide RNA tool (IDT). For each gene of interest, three crRNA sequences targeting early and late exons were selected (Table 2). One sequence targeted the *tyrosinase* (*tyr*) gene which does not impact M Cell regeneration. crRNA aliquots (10 nM each) were complexed with (1.2uM) of tracrRNA (IDT, #1072533) to create sgRNA. Guides were then complexed to 0.5 uM of cas-9 protein (IDT, #1081061) and injected into 1-cell stage embryos labeled with tg (Tol-056) within 20min after fertilization. Injection of three gRNA guides is reported to cause 90% biallelic mosaic KO of the targeted gene [82]. Presence of mutation was assayed using PCR flanking the predicted locations of the sites in injected animals.

### Statistics

Data was analyzed using GraphPad/Prism software. Specific statistical tests used are indicated in figure legends throughout the text.

---

**Supplemental Figure 1:**
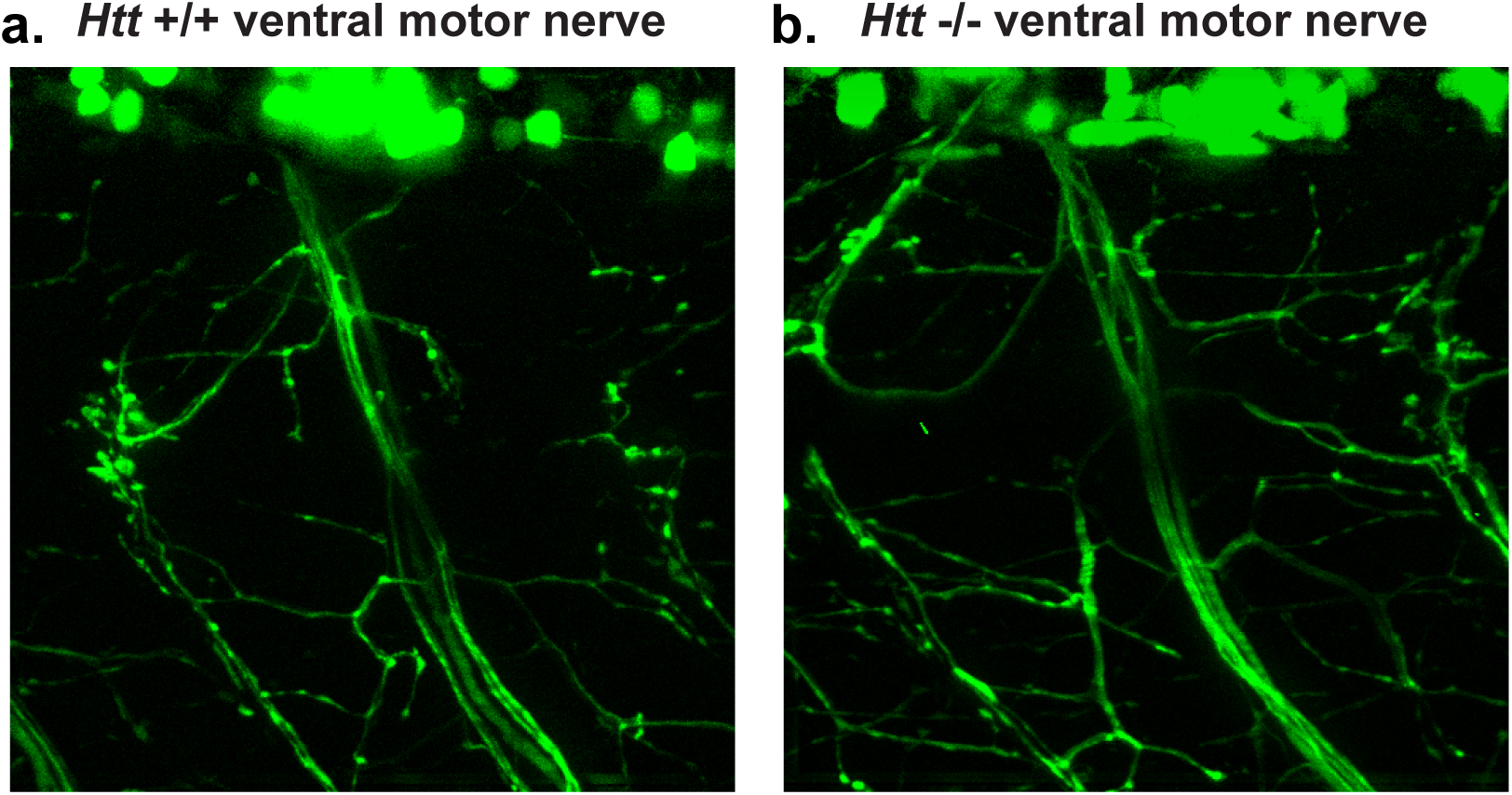
Ventral motor nerve development unaffected in *htt* mutants. (a-b) Images of non-transected ventral motor nerves *htt*+/+ and *htt*-/- samples (5dpf). htt-/- animals display no obvious deficits in motor nerve development.

**Supplemental Figure 2:**
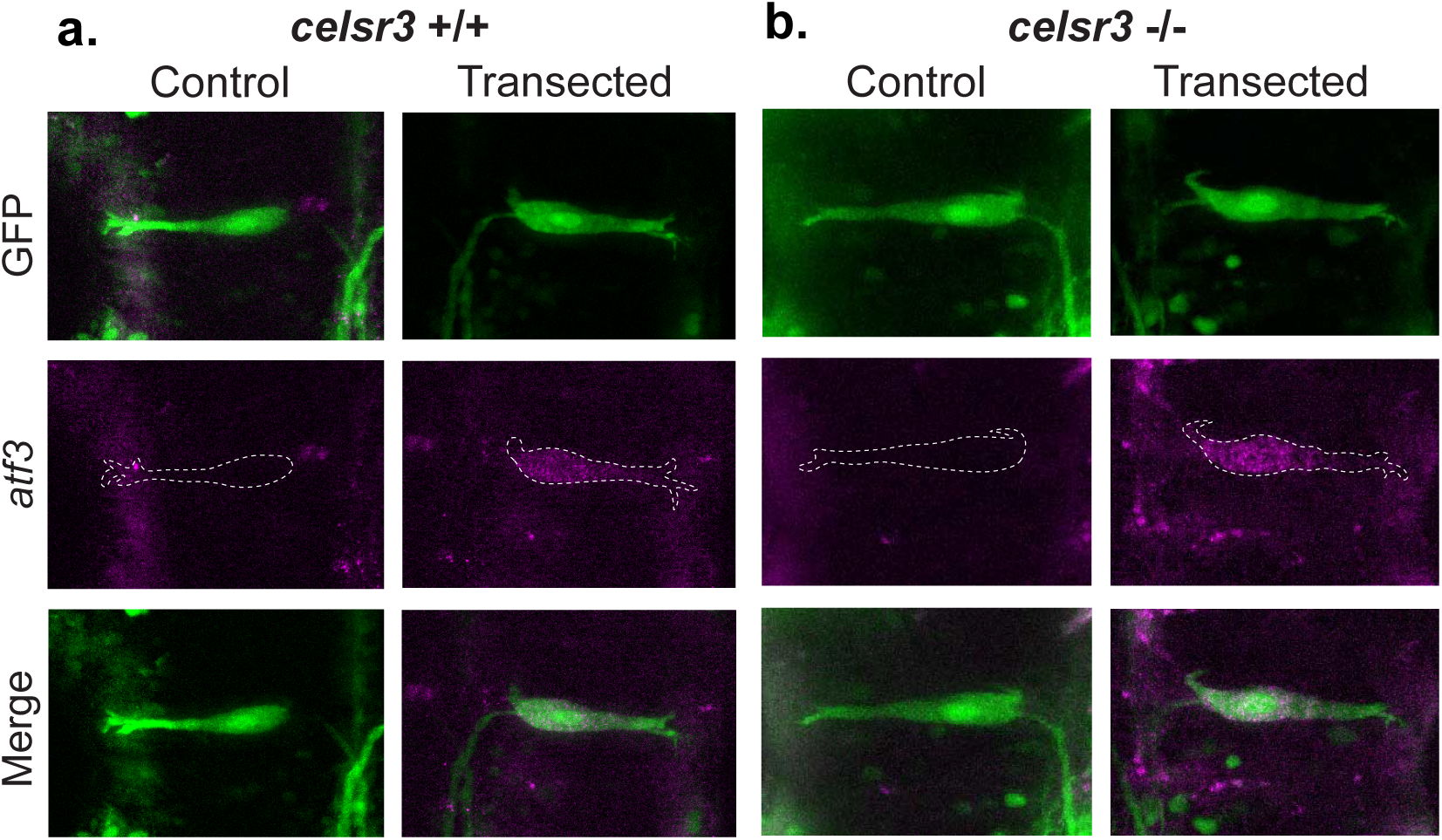
*celsr3* mutants upregulate *atf3* at 24 hpt. (a-b) Representative images of *atf3* upregulation in transected *celsr3* +/+ and *celsr3* -/- M-cells. *Celsr3* -/- animals display no obvious deficits in *atf3* upregulation.

**Supplemental Figure 3:**
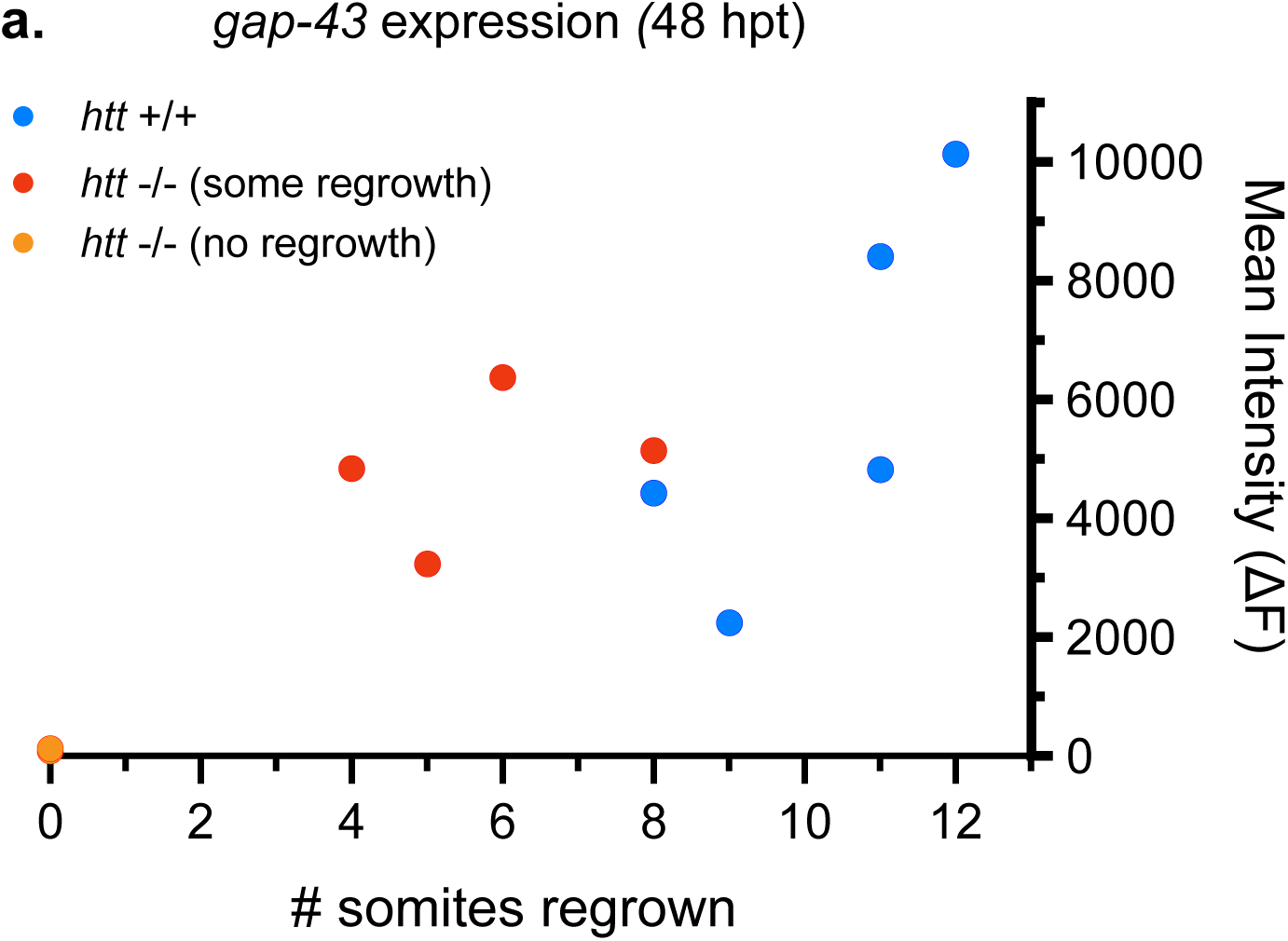
higher *gap-43* expression correlated with greater extent of regrowth in *htt* mutants. (a) Scatter plot of a subset of *htt* +/+ and *htt* -/- samples depicted in figure 5I. *gap43* expression at 48hpt within injured M-cells was plotted against the extent of regeneration at 48hpt within each sample. *htt* -/- samples with highly reduced *gap-43* expression at 48hpt also displayed no regrowth (n=2, orange) while *htt -/-* samples (n=4, red) displaying comparable *gap-43* expression to *htt* +/+ siblings (n=5, blue) displayed some regrowth.

## Movie Legends

Movie 1: Brief extension and retraction of axon tip in wildtype M-cell axon

Movie 2: Forward extension of axon tip in wildtype M-cell axon

Movie 3: Regenerating wildtype M-cell axon 4-24 hpt. Axon displays forward extension with little branching or misdirection.

Movie 4: Regenerating *htt^-/-^* M-cell axon 4-24 hpt. Axon displays delayed forward extension and excessive branching in injury site.

Movie 5: Regenerating *htt^-/-^* M-cell axon 4-24 hpt. Axon displays extended extension and retraction with no forward growth.

Movie 6: Regenerating wildtype M-cell axon 28-50 hpt. Axon displays rapid forward extension.

Movie 7: Regenerating *htt^-/-^* M-cell axon 28-50 hpt. Axon displays slow dynamics in forward extension.

